# Skin TDP-43 pathology as a candidate biomarker for predicting amyotrophic lateral sclerosis decades prior to motor symptom onset

**DOI:** 10.1101/2025.04.10.648122

**Authors:** Fergal M. Waldron, Tatiana Langerová, Aydan Rahmanova, Fiona L. Read, Holly Spence, Kristine Roberts, Angus D. Macleod, Samuel B. Pattle, Katie Hanna, Jenna M. Gregory

## Abstract

The recognition that disease-associated proteinopathies can manifest in peripheral organs outside the central nervous system preceding the onset of neurological symptoms, has transformed our understanding of Parkinson’s disease, in wide terms of pathogenesis, detection and diagnosis. For amyotrophic lateral sclerosis, non-motor symptoms, and non-central nervous system pathologies are gaining increased recognition but remain incompletely understood.

Here, using a TDP-43 RNA aptamer and a *Stathmin-2* cryptic exon transcript BaseScope^TM^ ISH probe, we identify widespread peripheral organ TDP-43 pathology prior to motor symptom onset in a discovery cohort of ante-mortem tissues from people who went on to develop ALS. Peripheral organs exhibiting both TDP-43 toxic gain- and loss-of function include muscle, lymph node, gallbladder, colon and with notably high incidence, skin. Given the accessibility of skin as a readily biopsiable tissue, representing a promising substrate for the detection of disease-associated proteinopathies and the development of minimally invasive biomarkers, we established an extended cohort of ante-mortem skin samples for TDP-43 pathology validation and further investigation. In skin biopsies taken during life from 17 individuals who went on to develop ALS we identify TDP-43 pathology from all 17 individuals in a wide distribution of anatomical sites, up to 26.5 years before ALS diagnosis – a presymptomatic period comparable to that observed for skin α-synucleinopathy in Parkinson’s disease. TDP-43 pathology was most abundant in skin biopsies from the back and shoulder, with sweat and sebaceous glands showing the highest involvement. TDP-43 pathology was also associated with structural changes.

As skin α-synucleinopathy has been established as a biomarker for both the detection of Parkinson’s disease and the differentiation of Parkinson’s disease from multiple system atrophy, we propose that skin TDP-43 likewise holds diagnostic and discrimination potential for diseases characterised by TDP-43 proteinopathy.

**Short Abstract:** Peripheral manifestations of neurodegenerative disease can precede neurological symptoms and serve as biomarkers, as shown by α-synuclein in the skin of individuals who later develop Parkinson’s disease. In amyotrophic lateral sclerosis (ALS), however, the distribution and diagnostic potential of peripheral TDP-43 pathology remain unclear. Using a TDP-43 RNA aptamer and a cryptic *STMN2* BaseScope™ probe, we examined ante-mortem tissues from individuals who later developed ALS. In a discovery cohort, we detected widespread pre-symptomatic TDP-43 pathology across multiple organs, with skin emerging as the most consistent site. We then validated these findings in a validation cohort comprising 17 individuals, all of whom exhibited TDP-43 pathology enriched in sweat glands and structural changes detectable up to 26.5 years before ALS diagnosis. These findings establish skin as a robust and accessible site of pre-symptomatic TDP-43 pathology, supporting its potential as a minimally invasive biomarker for early diagnosis and disease stratification in ALS.

**Summary:** Much like skin α-synucleinopathy has transformed biomarker development in Parkinson’s disease, this study identifies skin TDP-43 pathology as a promising early marker of ALS. The results open avenues for earlier diagnosis and stratification in a disease where intervention is most needed before symptoms appear.

**Graphical Abstract:** 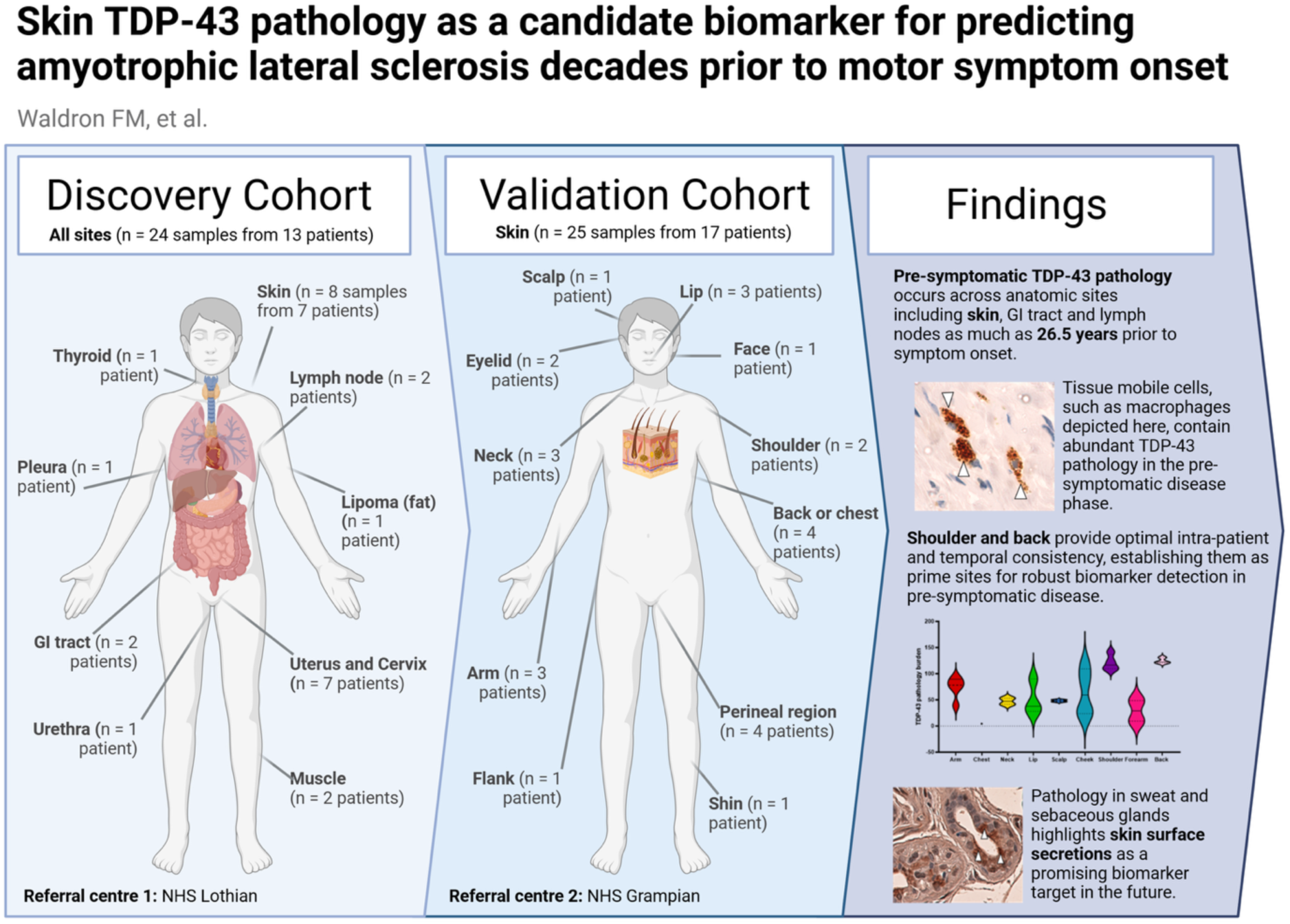

**Highlights:**

- Presymptomatic TDP-43 pathology occurs across a range on non-CNS peripheral organ systems including skin, gastrointestinal tract and lymph nodes prior to motor symptom onset in people who went on to develop ALS.
- In skin, presymptomatic TDP-43 pathology is associated with structural changes and can be detected up to 26.5 years prior to motor symptoms in ALS.
- As for Parkinson’s disease, shoulder and back represents optimal skin sampling sites for pre-symptomatic pathology in ALS.
- Sweat and sebaceous glands present with high levels of TDP-43 pathology, offering a promising biomarker target for early pathology detection.

**One Sentence Summary:** Using distinct biomarker discovery and validation ante-mortem tissue cohorts, we provide evidence of pre-symptomatic TDP-43 pathology across diverse non-CNS peripheral tissues, including skin decades before ALS symptom onset, highlighting skin TDP-43 pathology as a potential early biomarker for ALS and related TDP-43 proteinopathies

## Introduction

The recognition that neurodegenerative disease-associated proteinopathies can manifest in peripheral organs outside the central nervous system, preceding the onset of neurological symptoms, has transformed our understanding of Parkinson’s disease in terms of pathogenesis, detection, and diagnosis (*1–6*). This peripheral pathology perspective has reshaped the biomarker landscape, providing a path toward earlier, more accurate disease detection and the development of minimally invasive diagnostic tools. For example, skin α-synucleinopathy has been proposed a high potential biomarker for pre-symptomatic Parkinson’s disease (see review (*7*)), and has been shown to provide utility both for disease detection (*8, 9*), and also for distinguishing between Parkinsons’ disease and multiple system atrophy (*10, 11*). For amyotrophic lateral sclerosis (ALS), non-motor symptoms and non-central nervous system pathologies are gaining increased recognition but remain incompletely understood. Expanding the search for biomarkers beyond the central nervous system is therefore critical, not only for advancing disease biology but also for bridging the current gap in early detection strategies.

In ALS, the accumulation of pathological TDP-43 is the central hallmark (*12–14*). Once considered a purely motor disorder, ALS is now recognized to involve widespread non-motor and systemic manifestations, with pathological TDP-43 deposits detected years before symptom onset. These findings raise the possibility that peripheral tissues may provide early diagnostic or therapeutic entry points (*15–18*). These non-motor symptoms include cognitive symptoms of frontotemporal dementia (*19*), and systemic symptoms affecting a wider range of organ systems than the brain alone – indeed, extensive evidence of non-motor symptoms in non-CNS tissues for ALS is provided by pathological, patient reported, epidemiological, and clinical investigative studies, which extend to several organ systems and involvements including the gastrointestinal system (*20–35*), skin (*28, 36–49*), bladder (*27*), blood vessels (*40, 50*), lymph nodes (*28*), skeletal muscle (*51–53*) and myocardial muscle (*51–53*). Crucially, non-CNS pre-symptomatic TDP-43 pathologies have been shown in ALS by identifying toxic TDP-43 gain-of-function (*28, 53, 54*), or TDP-43 loss-of-function through splicing repression (*55*) prior to clinical symptoms, and sometimes over a decade before (*28*), indicating that early non-motor symptoms, and non-CNS tissues along with fluid biomarkers, provide potential for early diagnosis of ALS.

Despite this promise, progress toward peripheral TDP-43 biomarkers has been hindered by limitations of classical antibody-based detection methods, which lack either specificity (total TDP-43 antibodies) or sensitivity (pTDP-43 antibodies) for TDP-43 pathology (*56*). We recently developed an RNA aptamer–based approach (TDP-43^APT^) that markedly improves detection of pathological TDP-43 aggregates (*14, 57, 58*). When combined with *in situ* hybridization (ISH) probes for cryptic *STMN2* exons, this strategy simultaneously captures gain- and loss-of-function consequences of TDP-43 dysregulation, events that conventional antibody methods fail to reveal (*14*). Together, these new tools facilitate the detection of a wider range of pathological events compared to classical antibody approaches alone, providing crucial insights in ALS including early nuclear TDP-43 aggregation preceding cytoplasmic mislocalisation (*59*), and evidence of TDP-43 pathology in SOD1-ALS (*59*).

Here, we take advantage of the improved specificity and sensitivity of this dual aptamer/ISH approach, to probe a discovery *ante-mortem* tissue cohort to establish the burden of systemic, non-CNS, TDP-43 pathology with greater sensitivity and specificity, and to assess whether TDP-43 loss-of-function events are detectable in these non-CNS tissues. Our goal was to identify accessible sites for pre-symptomatic diagnosis and trial biomarker development, while shedding light on mechanisms that may underpin the diverse non-motor manifestations of ALS. Having identified skin as a reproducible site to detect pathology we then established an extended validation cohort of ante-mortem skin samples comprising 17 individuals from a second referral centre who also later went on to develop ALS. Using this extended validation cohort, we were able to test the reproducibility of skin as a peripheral biomarker site and to assess the gap in knowledge surrounding the temporal and spatial distribution of peripheral TDP-43 pathology and its association with structural alterations within this tissue in the pre-symptomatic disease phase. By integrating both a discovery and an independent validation cohort, our study aims to directly address key knowledge gaps pertaining to peripheral pre-symptomatic disease states and to establish a robust framework for advancing peripheral TDP-43 pathology as a clinically meaningful biomarker in ALS.

## Results

In this study, with the aim of identifying novel potential biomarker modalities for early detection of ALS and TDP-43 proteinopathies, we investigate the extent of pre-symptomatic TDP-43 pathology in non-CNS peripheral organ systems from tissue biopsies taken during life from people who went on to develop ALS. Employing recently developed molecular pathology detection tools (TDP-43 RNA aptamer, and *STMN-2* cryptic exon BaseScope^TM^ in situ hybridisation probe) with greater sensitivity and specificity than current antibody approaches, we uncover extensive organ and cell type involvement in a biomarker discovery cohort of ante-mortem peripheral tissues from 8 organ systems, and identify skin TDP-43 pathology as a candidate biomarker showing a high instance of pre-symptomatic pathological TDP-43. In a separate biomarker validation cohort of ante-mortem skin biopsies taken from people who went on to develop ALS, we establish skin TDP-43 pathology as a potential pre-symptomatic ALS biomarker, occurring at very high incidences, across a wide-range of anatomical sites, from up to 26.5 years prior to ALS symptom onset.

### Results section 1: Biomarker discovery cohort (peripheral organ systems)

The biomarker discovery cohort consisted of non-CNS peripheral tissue biopsies taken ante-mortem from people (Lothians, Scotland, UK) who went on to develop ALS, and was sourced from the NHS Lothian Tissue Biorepository in Edinburgh, UK.

#### Pre-symptomatic TDP-43 pathology spans multiple non-CNS peripheral organ systems

In our ALS pre-symptomatic peripheral tissue TDP-43 pathology discovery cohort, we examined tissues from eight organ systems across 13 individuals for evidence of TDP-43 pathology using a TDP-43 RNA aptamer and a *STMN-2* cryptic exon BaseScope™ in situ hybridisation probe. We identified pre-symptomatic TDP-43 pathology in several peripheral organs and tissues, including skin, muscle, colon, gallbladder, lymph node (Figure 1A; Table 1). Cell types affected include sebocytes, endothelial cells, peripheral neuronal cells, dendritic cells, chondrocytes - all cells with a shared cell linage from the neural crest (Figure 1A). In our discovery cohort, TDP-43 pathology in non-CNS peripheral tissues was detectable up to 2–14 years before ALS symptom onset in lymph nodes, 1–11 years in skin, and 1–2 years in the gastrointestinal tract (Figure 1B). In contrast, we found no evidence of TDP-43 pathology in other peripheral tissues such as pleura, bladder, cervix, urethra, uterus, thyroid, and from a lipoma (Figure 1A; Table).

**Figure 1.**
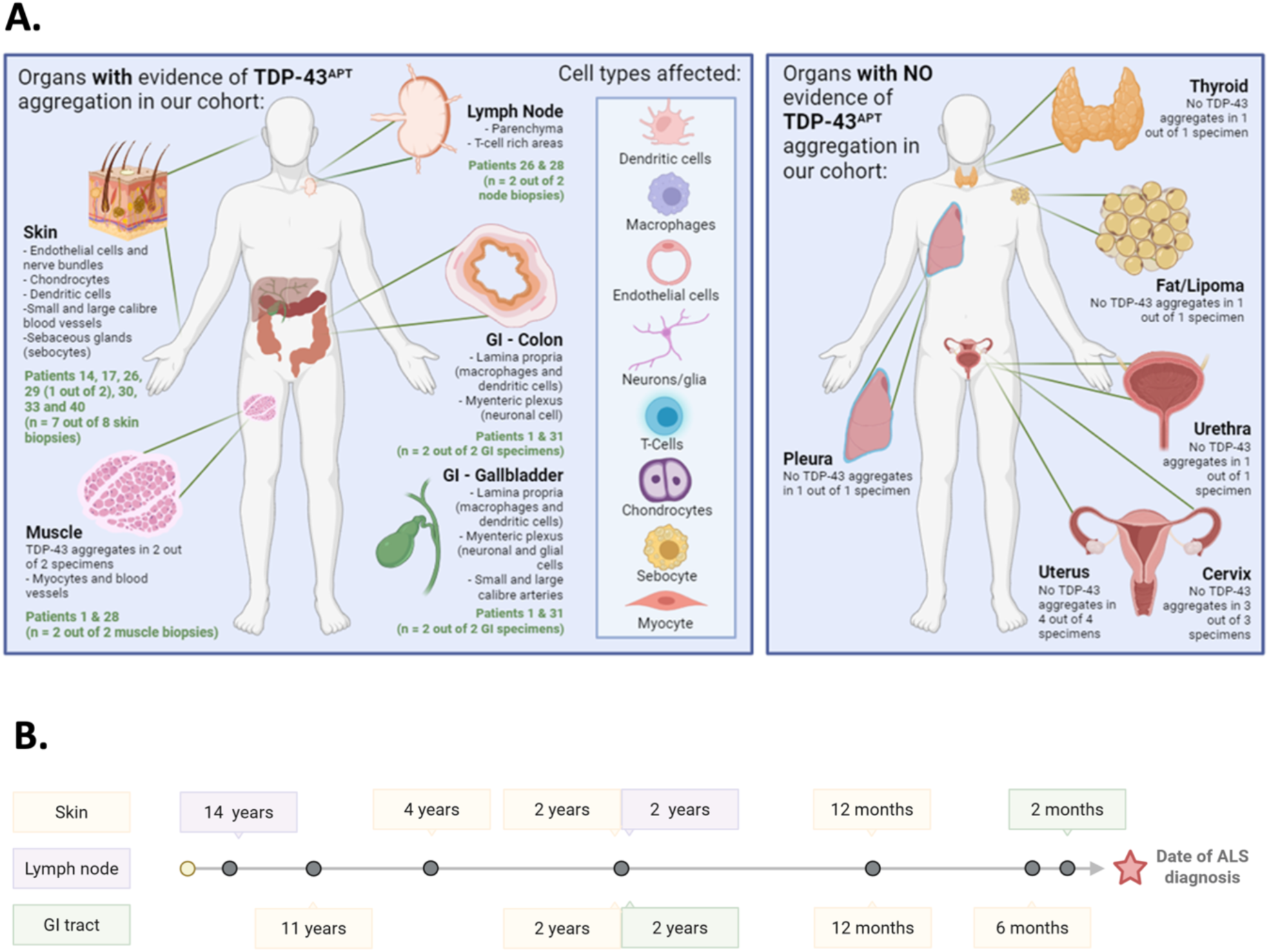
Improved sensitivity of TDP-43^APT^ facilitates detection of TDP-43 pathology in discovery cohort of ALS pre-symptomatic peripheral tissues. **A.** Schematic demonstrating organs with evidence of TDP-43 pathology, detected (left), and those with no evidence of TDP-43 pathology (right). Cell types affected by TDP-43 pathology are also detailed in the inset box in the left panel. **B.** Schematic demonstrating range of time from *ante-mortem* sampling to symptom onset across organs that show evidence of TDP-43 pathology.

**Table 1.**
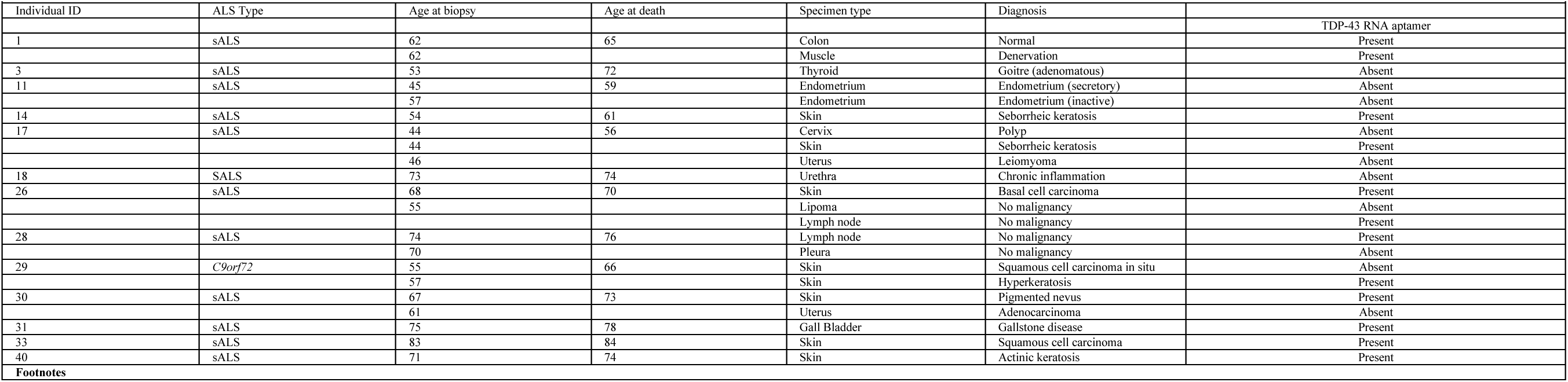
Widespread pre-symptomatic TDP-43 pathology revealed from biomarker discovery cohort of ante-mortem peripheral tissue biopsies taken from people who went on to develop ALS. Summary table of clinicopathological information for the biomarker discovery cohort (n = 13 individuals), where non-CNS peripheral tissues sampled prior to ALS motor symptom onset were screened for TDP-43 pathology. Each row lists the individual ID, ALS subtype, age at biopsy, age at death, biopsy specimen type, contemporaneous pathological diagnosis, along with the presence or absence of TDP-43 pathology. For several individuals, multiple tissue samples collected at different times or from distinct anatomical sites are presented.

By combining TDP-43 RNA aptamer and *STMN-2* cryptic exon BaseScope™ in situ hybridisation probe approaches, we achieved improved sensitivity for detecting TDP-43 pathology in both skin and lymph node compared with previously used antibody-based approaches, and we were also able to reveal TDP-43 pathology in muscle that had not been detected with TDP-43 antibodies (see emental Table 1). Specifically, we demonstrate: (i) in skin, markedly improved sensitivity for detecting TDP-43 pathology compared with antibody-based methods (pathology detected in 7/8 biopsies from 7 patients versus 2/8 biopsies from the same number of patients previously; (ii) in lymph nodes, similarly enhanced detection sensitivity (2/2 biopsies from 2 individuals versus 1/2 previously); and (iii) in muscle, we reveal the presence of TDP-43 pathology that was not detectable using antibodies is revealed here.

Organs such as the gallbladder and colon, which were previously reported to contain TDP-43 aggregates detected by antibody-based methods (see Supplemental Table 1), also exhibited detectable gain- and loss-of-function TDP-43 pathology in the present study. Notably, we were unable to detect TDP-43 pathology in pleura, bladder, cervix, urethra, uterus, thyroid, and from a lipoma biopsy (Figure 1A; Table 1). The number of individuals and samples for these organ systems was low however, therefore further work to investigate involvement of these under-represented organs may be warranted.

#### Skin shows a high incidence of pre-symptomatic TDP-43 pathology amongst non-CNS tissues

In our TDP-43 pre-symptomatic peripheral pathology discovery cohort, seven individuals had undergone skin biopsies prior to the onset of ALS motor symptoms - Pre-symptomatic TDP-43^APT^ pathology was identified from skin in all seven individuals, ranging from 0.5-11 years (median 2 years; Q1-Q3 = 1-4 years; Figure 1B; Table 1) prior to symptom onset.

TDP-43 pathology was observed most frequently within dendritic cells of the dermis, peripheral nerve bundles and small dermal blood vessels (Figure 2A). Notably, one individual had two skin biopsies taken during life prior to ALS symptoms, the first without detectable TDP-43 pathology but the second with detected TDP-43 pathology representing a potential phenoconversion event (Figure 2B) prior to symptom onset. This individual, who had a *C9orf72* mutation, had two biopsies taken from the same site (case 29; Supplementary Figure 1), the first of which was taken 24 months prior to symptom onset, with the second biopsy taken within 12 months of diagnosis (Figure 1B). The first biopsy taken 24 months prior to ALS symptom onset showed no evidence of TDP-43 pathology, however, the second biopsy showed florid TDP-43 pathology within the dendritic cells of the dermis, the sebocytes within the sebaceous glands and within the peripheral nerve bundles surrounding the sebaceous glands. The peripheral nerve pathology was confirmed using *STMN-2* cryptic exon *in situ* hybridisation probes detecting neuron-specific TDP-43 loss of function (Figure 2C).

**Figure 2.**
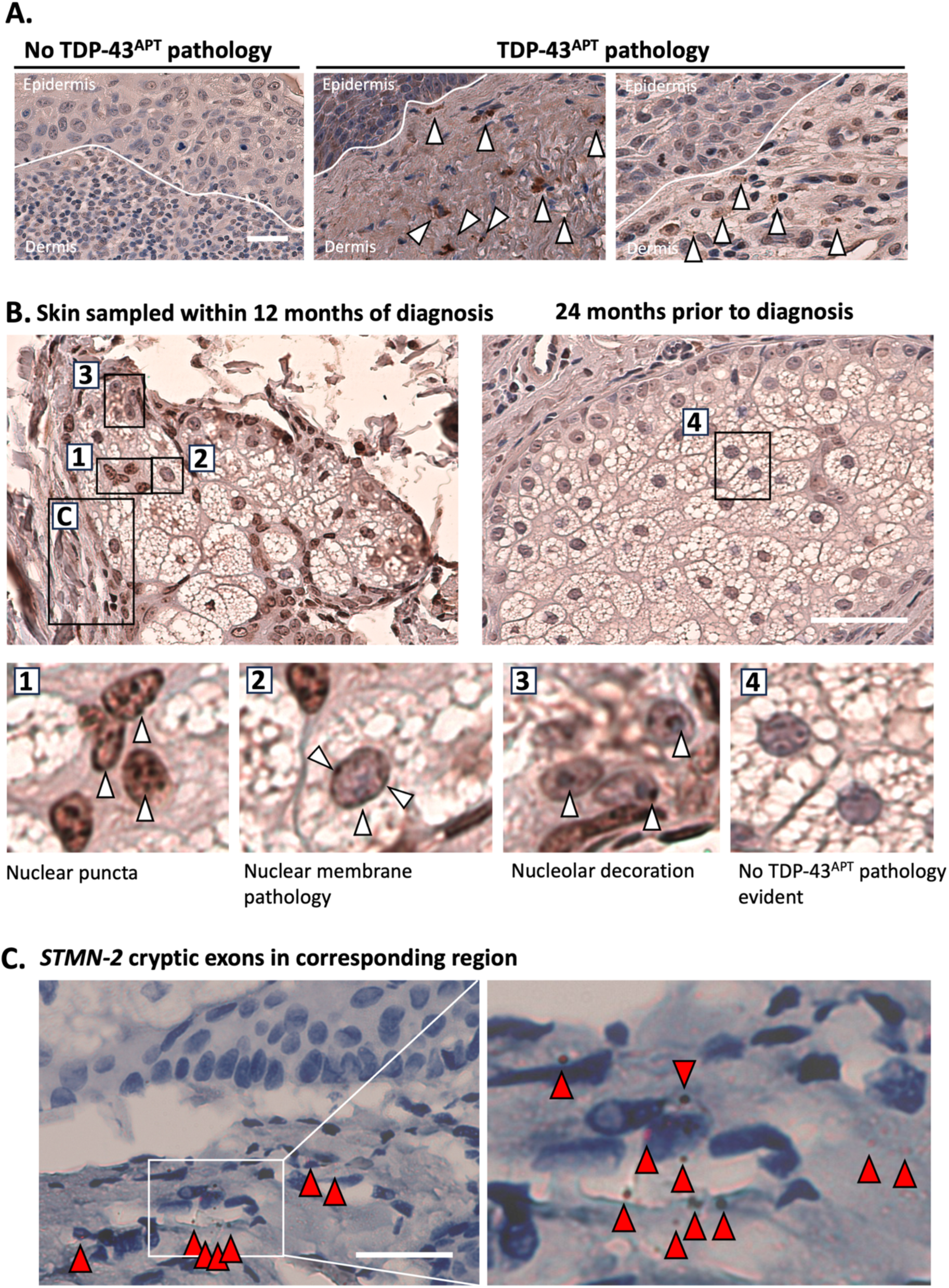
TDP-43^APT^ consistently detects TDP-43 pathology in skin biopsies with improved sensitivity compared to antibody approaches in pre-symptomatic ALS. **A.** Photomicrographs taken at 40x magnification stained with TDP-43^APT^ with DAB chromogen and counterstained with haematoxylin. Images show examples of a single case that had no evidence of TDP-43^APT^ pathology (left) and images from two other cases that showed evidence of TDP-43^APT^ pathology within the dermis (white arrowheads) of individuals who later went on to develop ALS. Scale bar = 20 μm. **B**. Images of skin biopsies from one individual who had a *C9orf72* mutation. The image on the left, taken the same year as symptom onset, shows evidence of TDP-43^APT^ pathology (white arrowheads) in the nuclei and cytoplasm of sebocytes of the sebaceous gland within the dermis (1–3). The image shows the same site biopsied 36 months prior to diagnosis, with no evidence of TDP-43^APT^ pathology (4). Scale bar = 40 μm **C**. Image taken from area highlighted in (B) showing the presence of *STMN-2* cryptic exons (indicated by red arrowheads) in the peripheral nerve bundle adjacent to the affected sebaceous gland. Scale bar = 50 μm.

#### Muscle TDP-43 pathology in myocytes and surrounding neurovascular bundles

Whilst pTDP-43 antibody staining failed to identify TDP-43 aggregates in muscle (Figure 3A), a serial section from the same muscle biopsy stained with TDP-43^APT^ revealed TDP-43 puncta within (i) the cytoplasm of the myocytes (sarcoplasm) and (ii) the myocyte nuclei (myonuclei) which are dispersed along the inner surface of the sarcolemma at the periphery of each fiber (Figure 3A, B). Furthermore, TDP-43 pathology was observed within the neurons of the peripheral nerve bundles running between fascicles (Figure 3C), confirmed by *in situ* hybridisation detection of *STMN-2* (Figure 3D, Supplementary Figure 2). Sebocytes and myocytes represent two more non-CNS cells types with identified TDP-43 pathology, facilitated here by the increased sensitivity of our TDP-43 aptamer and STMN-2 BaseScope^TM^ probe, to add to those where pTDP-43 had been identified using classical antibody approaches (i.e. dendritic cells, macrophages, endothelial cells, T-cells and chondrocytes; (*28*)

**Figure 3.**
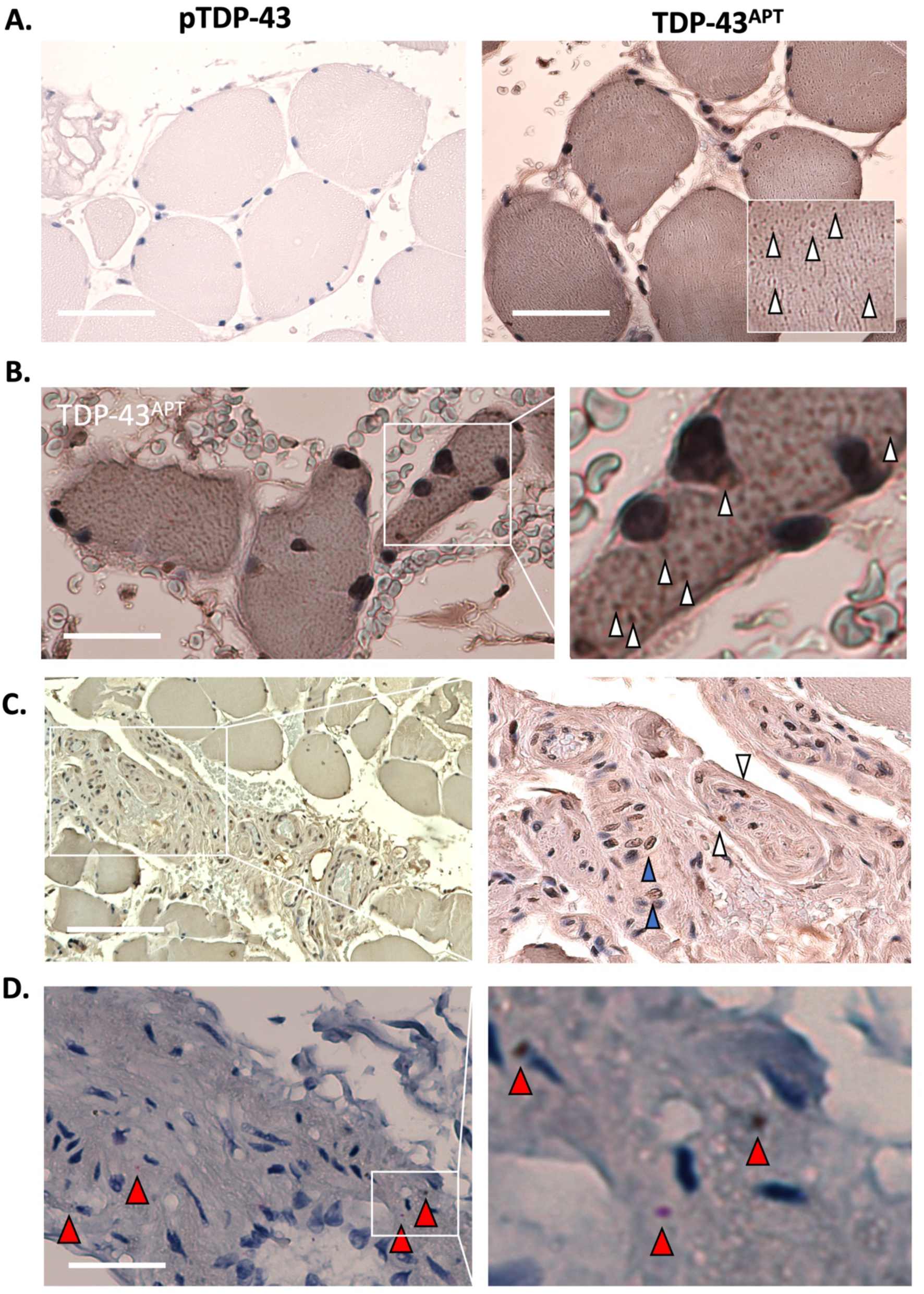
Muscle biopsy reveals TDP-43 pathology in myocytes and surrounding neurovascular bundles in pre-symptomatic ALS. **A.** Photomicrographs taken at 40x magnification stained with pTDP-43 antibody (left image) and TDP-43^APT^ (right image) with DAB chromogen and counterstained with haematoxylin. Scale bar = 50 μm. Images demonstrate lack of immunoreactivity for pTDP-43 antibody, but strong immunoreactivity for TDP-43^APT^ staining. **B**. Photomicrographs taken at 20x magnification stained with TDP-43^APT^ (left image) with 4x optical zoom in right image. These images are taken from the second muscle biopsy indicating that TDP-43^APT^ pathology can be seen in both individuals sampled within this cohort. White arrowheads indicate TDP-43^APT^ pathology; scale bar = 50 μm. **C**. Photomicrographs taken at 20x magnification stained with TDP-43^APT^ (left image) with optical zoom in right image. Scale bar = 75 μm. The image demonstrates TDP-43^APT^ pathology within the neurovascular bundle in between fascicles. White arrowheads indicate TDP-43^APT^ pathology within the peripheral nerve, and blue arrowheads indicate pathology within the endothelial cells of the adjacent blood vessels. **D**. The same region imaged in (C), except stained with *in situ* hybridisation probes directed at *STMN-2* cryptic exon, which is only seen in the context of TDP-43 loss-of-function. Here TDP-43 loss-of-function can be visualised within the peripheral nerve bundle (red arrowheads indicate individual mRNA containing the *STMN-2* cryptic exon. Scale bar = 50 μm.

#### Blood vessel involvement in pre-symptomatic ALS

A contingent of small blood vessels (mostly arterioles, venules and small veins) was present in all peripheral biopsies in this cohort. Organs with demonstratable TDP-43 pathology, i.e., skin, muscle, lymph nodes, colon and gall bladder, contained both TDP-43 positive and negative blood vessels (Figure 4).

**Figure 4.**
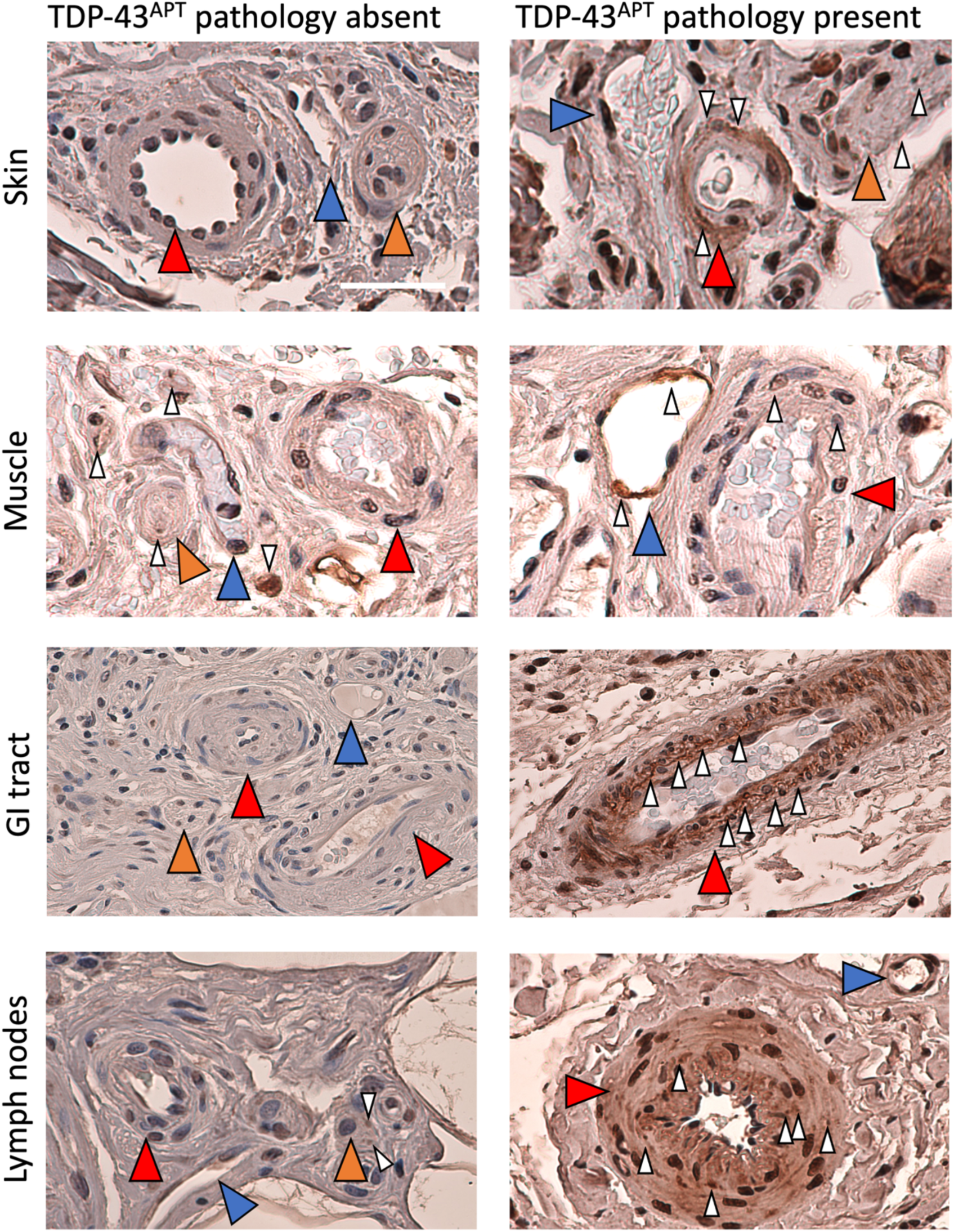
Blood vessel involvement by TDP-43 pathology in pre-symptomatic ALS. Photomicrographs taken at 20x magnification stained with TDP-43^APT^ with DAB chromogen and counterstained with haematoxylin. Images demonstrate examples of neurovascular bundles with and without TDP-43^APT^ pathology from each of the affected organ systems. Scale bar = 50 μm. Red arrowheads indicate arteries, blue arrowheads indicate veins, orange arrowheads indicate peripheral nerves. White arrowheads indicate TDP-43^APT^ pathology.

#### Aggregate-laden macrophages present across non-CNS tissues including gastrointestinal tract, skin and lymph nodes

Amongst non-CNS tissues examined here, we note the presence of morphologically abnormal macrophages. Aberrant morphologies include aggregate-laden macrophages, staining positively for TDP-43 pathology, and macrophages full of large vacuoles indicative of lysosomal dysfunction. We observed such aberrant macrophage morphologies in gastrointestinal tract, skin and lymph node specimens (Figure 5) – all of which were positive for pathological TDP-43.

**Figure 5.**
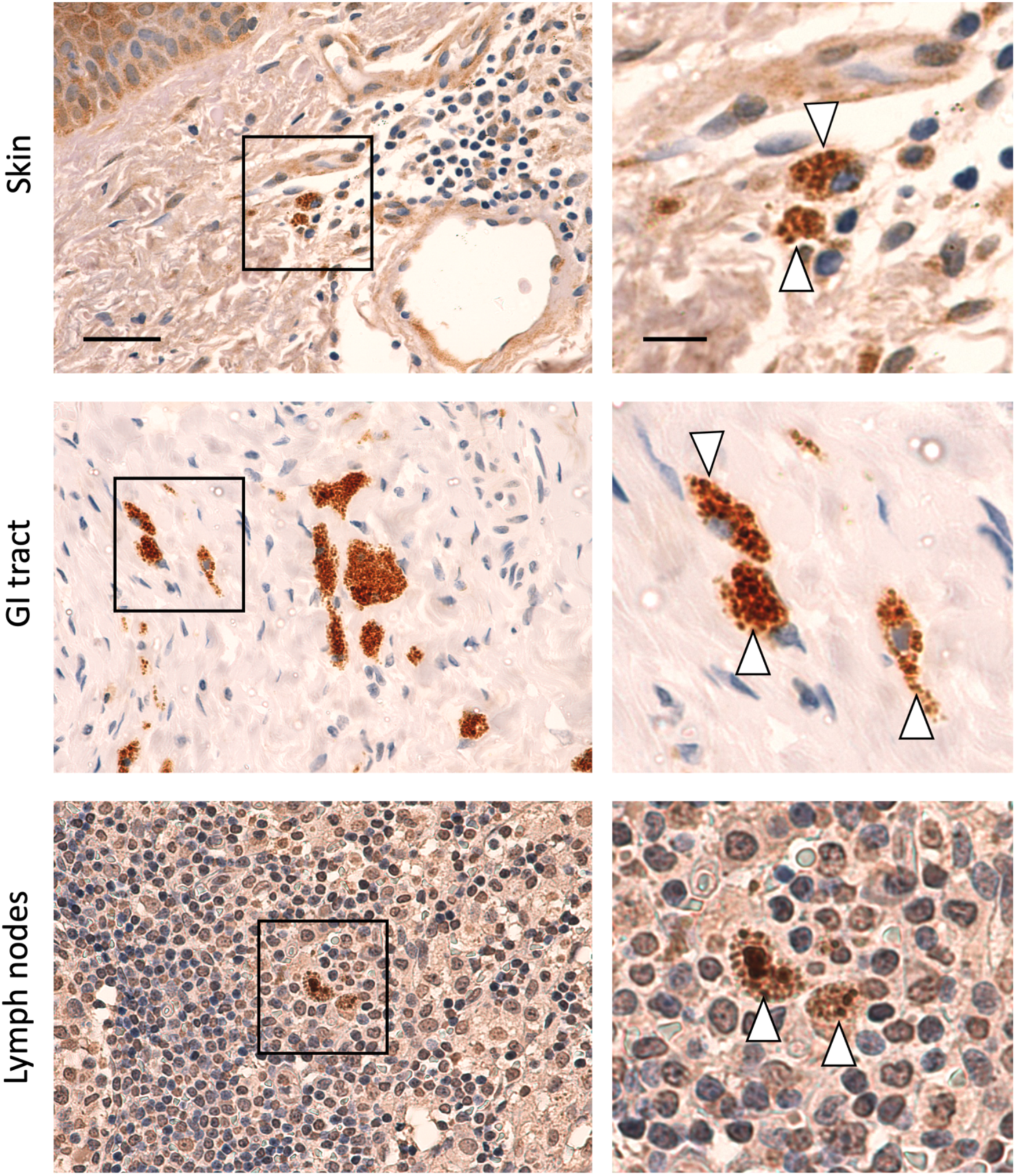
Aggregate- and vacuole-laden macrophages observed across non-CNS tissues in pre-symptomatic ALS. Photomicrographs taken at 40x magnification stained with TDP-43^APT^ with DAB chromogen and counterstained with haematoxylin. Scale bar = 50 μm in the left panel and 20 μm in the right panel. Black box in left panel represents area of optical zoom in right panel. Amongst non-CNS tissues we observed the presence of abnormal macrophage morphologies (white arrowhead) including aggregate- and vacuole-laden macrophages, in gastrointestinal tract, skin, and lymph node, consistent with histological evidence of lysosomal dysfunction.

### Results section 2: Biomarker validation cohort (skin)

The biomarker validation cohort consisted of skin biopsies taken ante-mortem from people (Grampian, Scotland, UK) who went on to develop ALS, and was sourced from the NHS Grampian Tissue Biorepository in Aberdeen, UK.

#### Skin TDP-43 pathology is a candidate biomarker for early ALS detection

Having detected pre-symptomatic skin TDP-43 pathology, ranging from 0.5-11 years before motor symptom onset, from all 7 individuals in our discovery cohort of peripheral tissue biopsies, we established an expanded biomarker validation cohort for skin to further investigate the prevalence and chronological range of TDP-43 pathology detectable in skin prior to motor symptom onset in people who went on the develop ALS.

In this separate, expanded validation cohort of 25 skin biopsies from 17 individuals across a wide range of anatomical sites (Figure 6A), TDP-43 pathology was present in all 25 biopsies from all 17 individuals, from 213 days up to 26.5 years before motor symptom onset (Figure 6B; Table 2. The median time between pre-symptomatic skin TDP-43 pathology and motor symptom onset for TDP-43 positive biopsies in our skin validation cohort was 10.59 years (Q1-Q3; 5.29-18.45) or 9517 days.

**Figure 6.**
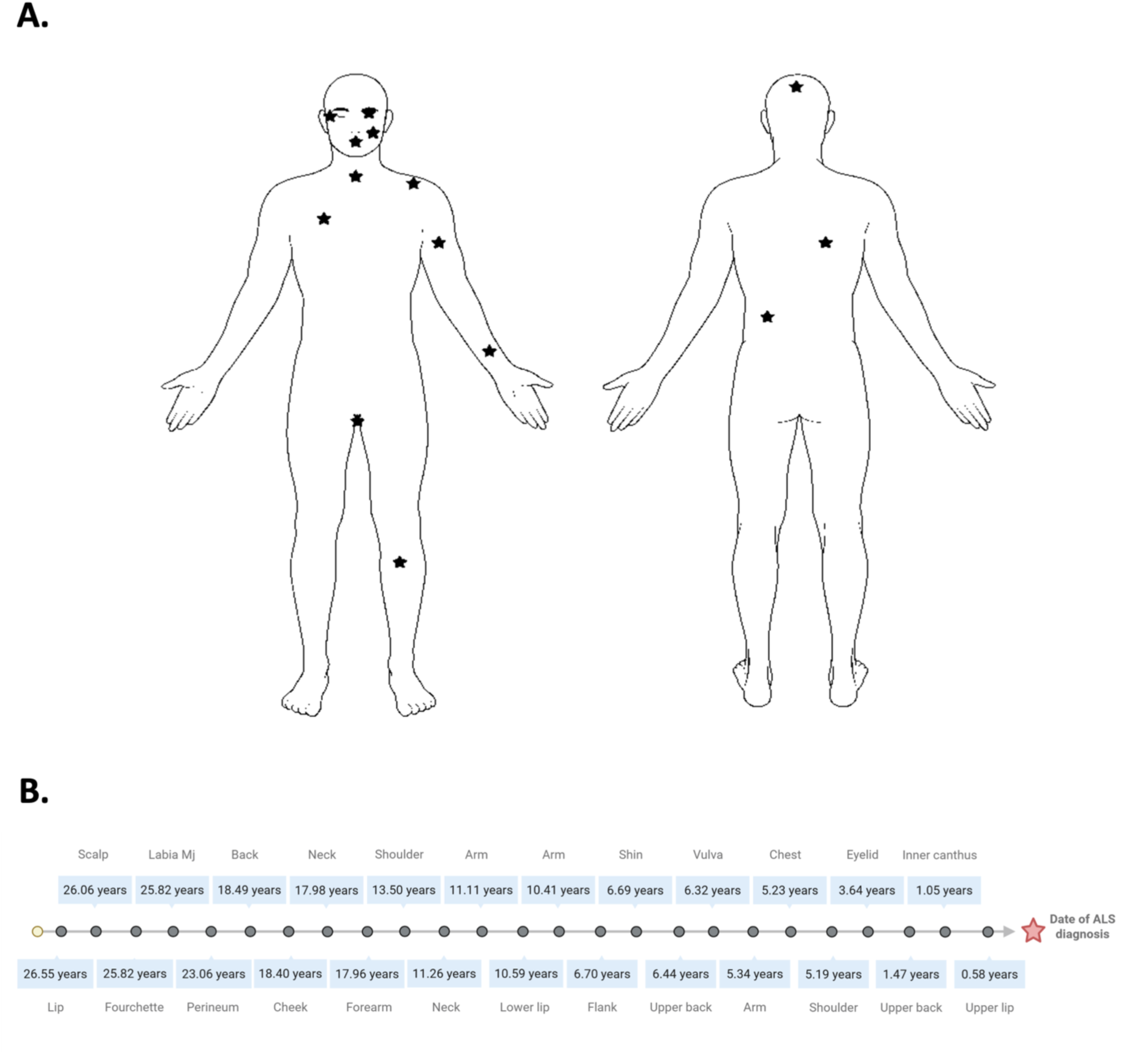
Widespread presymptomatic skin TDP-43 pathology revealed from extended validation cohort of skin biopsies up to 26.5 years prior to motor symptom onset. **A.** Anatomical illustrations of the wide-range of anatomical sites from which TDP-43 pathology was detected from skin. **B.** Timeline representing years prior to ALS diagnosis that all 25 TDP-43 pathology positive skin biopsies had been taken, ranging from 0.58 years to 26.57 years, with anatomical sites of biopsies.

**Table 2.**
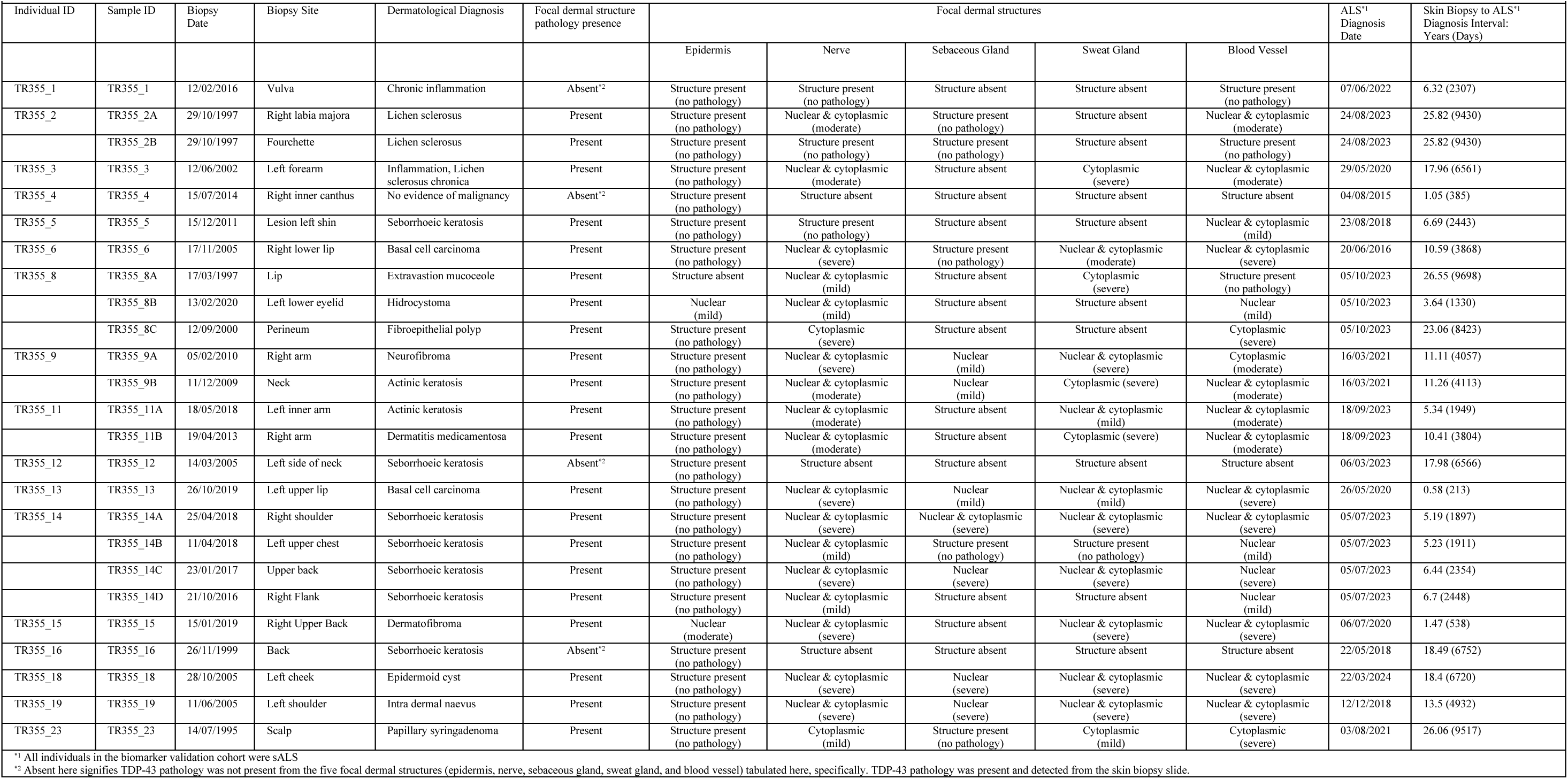
Widespread pre-symptomatic skin TDP-43 pathology revealed from biomarker validation cohort of ante-mortem skin biopsies taken from people who went on to develop ALS. Summary table of clinicopathological information for the biomarker validation cohort (n = 25 individuals), where skin biopsies sampled prior to ALS motor symptom onset were screened for TDP-43 pathology. Each row reports the individual ID, sample ID, biopsy date, biopsy site, dermatological diagnosis, dermatological diagnosis, along with histological features of dermal structures, and the presence, localization (nuclear and/or cytoplasmic), and semi-quantitative severity of TDP-43 pathology in distinct skin compartments (epidermis, nerves, sebaceous glands, sweat glands, and dermal blood vessels). The table also provides the ALS diagnosis date and the interval between the skin biopsy and the ALS diagnosis (expressed in years and days). Multiple biopsies from the same individual, sometimes at different anatomical sites or times, are reported separately to illustrate intra-individual heterogeneity of early TDP-43 pathology.

Skin TDP-43 pathology detected from a wide range of anatomical sites including head and neck, trunk, perineal area and the upper and lower limb (Figure 6A), but crucially TDP-43 pathology was detected universally across all skin biopsies, indicating a lack of anatomical predisposition for TDP-43 pathology in skin.

It should be noted that from instances where skin biopsies were available from multiple sites from the same individual. Indeed, for 4 individuals in our validation cohort, TDP-43 pathology-positive skin biopsies were available from multiple sites from the same individual – for these individuals, multiple site biopsies were taken either on the same day (TR355_2 Katie), 8 weeks apart (TR355_9 Katie), 1.5 years apart (TR355_14) or in one individual (TR355_8) 19.5 years apart (see Table 2). Specifically, individual TR355_2 hasTDP-43 pathology from 2 vulval sites (labia majora and labia minora) biopsied on the same day, over 25.8 years prior to symptom onset (PSO). Individual TR355_9 showed TDP-43 pathology from 2 sites biopsied 56 days apart in order of neck (11.3 years PSO) and them arm (11.1 years PSO). Individual TR355_14 had TDP-43 pathology from 4 sites biopsied in order of right flank (6.7 years PSO), left upper back (6.4 years PSO), left upper chest (5.2 years PSO), and right shoulder (5.2 years PSO). Remarkably, individual TR355_8 had TDP-43 pathology from 3 sites biopsied in order of lip (26.5 years PSO), perineum (23.1 years PSO) and lower eyelid (3.6 years PSO) (see Table 2).

#### TDP-43 skin pathology predominantly in peripheral nerves, blood vessels, and adnexal structures

TDP-43 pathology was predominantly observed in peripheral nerves, blood vessels, and adnexal structures including sweat and sebaceous glands (Figure 7A). Each of these four dermal structures, along with epidermis, was manually graded, as described previously (*14*), with the extent and pattern of TDP-43^APT^ staining recorded (Table 2). Not all structures were present in every biopsy; where absent, pathology was instead evident in other cell types or structures such as macrophages and dendritic cells.

**Figure 7.**
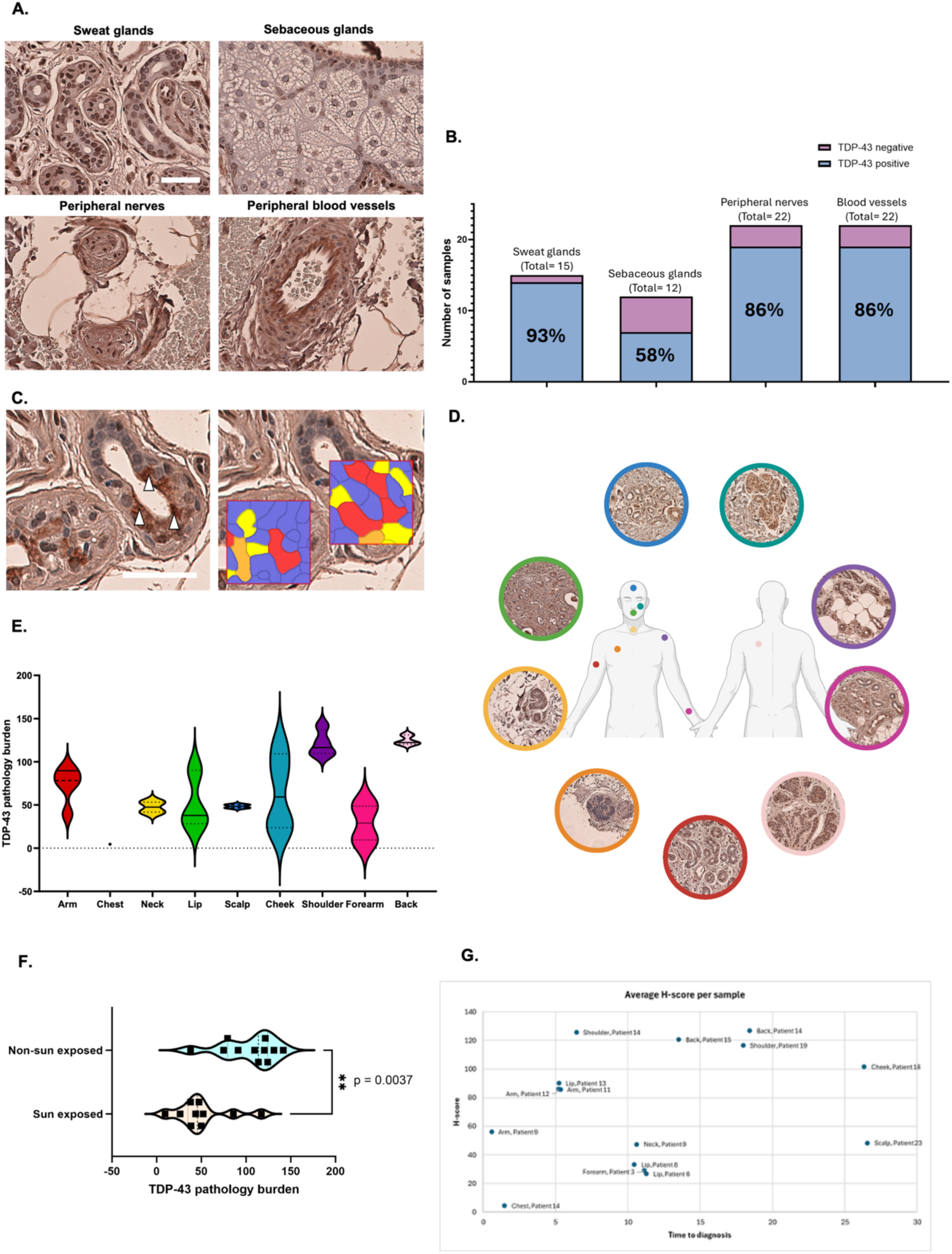
Sweat and sebaceous glands show highest TDP-43 pathology. TDP-43 pathology burden across peripheral neurovascular bundles and adnexal structural and its anatomical and temporal variation in sweat glands is illustrated here in the extended validation cohort. **A.** Histological images demonstrate the predominant location of TDP-43^APT^ pathology in this extended presymptomatic cohort. Scale bar = 50 μm. **B.** Proportion of dermal structures positive for TDP-43 pathology. Each bar represents the total number of biopsies in which sweat glands, sebaceous glands, peripheral nerves, and blood vessels were present. Bars show the proportion of these structures with detectable TDP-43 pathology (blue) versus those without pathology (pink). Sweat glands were TDP-43 positive in 93% of cases, sebaceous glands in 58%, and both peripheral nerves and blood vessels in 86%. **C.** Histology images illustrating the methodology of TDP-43 pathology burden scoring in sweat glands by superpixels-based segmentation. This method groups pixel similarity between different cellular populations and provides information pertaining to DAB intensity, defined as a ‘H-score’. Scale bar = 50 μm. **D.** Anatomical illustration of sites where skin biopsies included sweat glands, with representative histological images. These regions included the scalp, cheek, shoulder, forearm, back, arm, chest, neck and lip. **E.** Violin plots for TDP-43 pathology burden across anatomical regions for which skin biopsies contain sweat glands. TDP-43 pathology burden was greatest in the sweat glands of the shoulder and back and lowest in that of the chest Anatomical regions are colour-coded in correspondence with Figure 7D., previously. **F.** Violin plots for TDP-43 pathology burden for sun exposed (forearm, lip, neck, scalp and cheek), and non-sun exposed (arm, shoulder and back), anatomical sites with representative p-value (p=0.0037; Mann Whitney-U test) showing that pathology burden was significantly greater in non-sun exposed skin regions. **G.** TDP-43 pathology burden (H-score) of sweat glands from a range of anatomical sites was plotted against time to ALS diagnosis. Heterogeneous trajectories were observed, with some regions (e.g., back, shoulder) consistently showing high burden, while others (e.g., chest, lip, arm) varied between patients and proximity to symptom onset.

**Figure 8.**
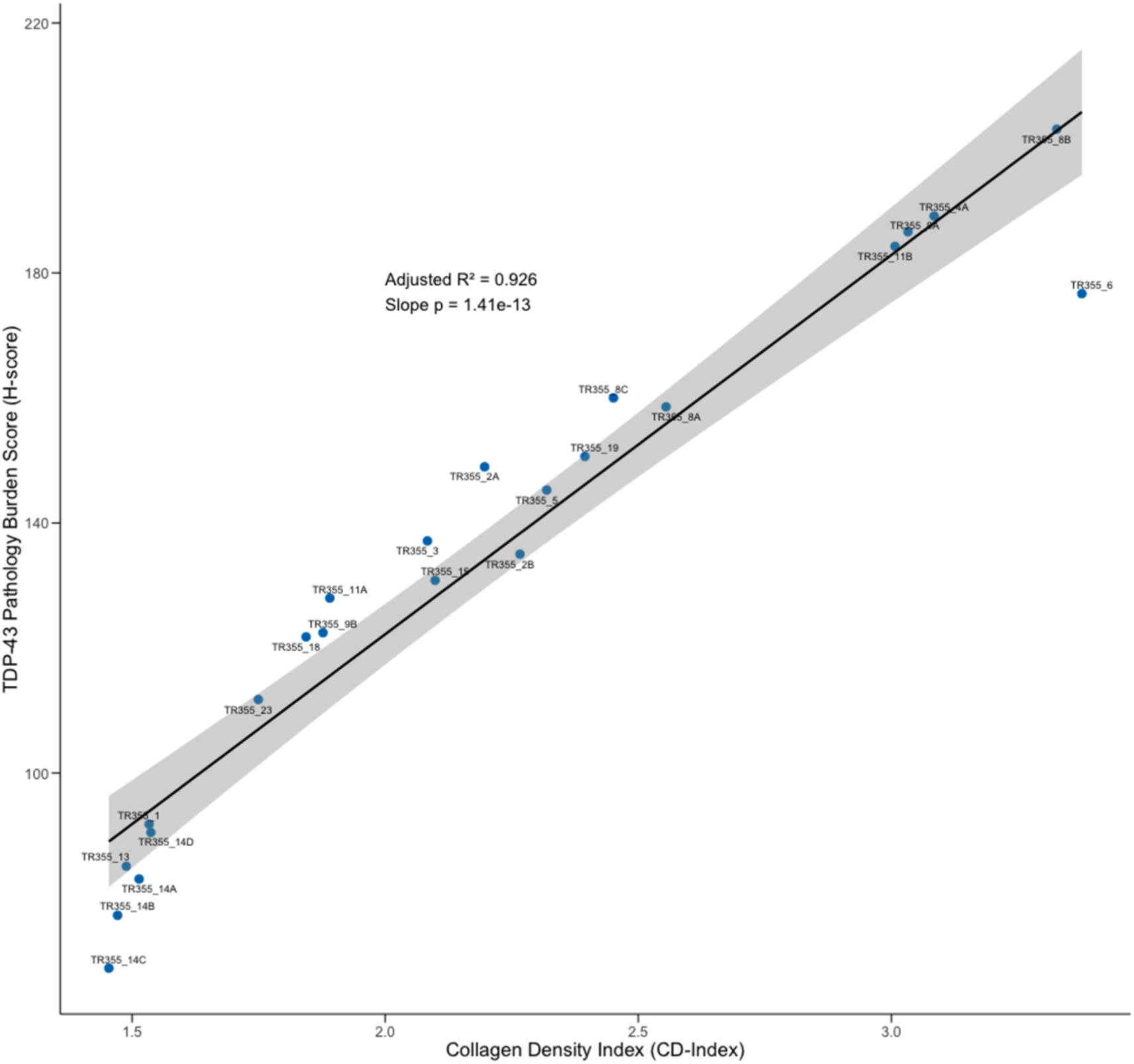
Skin collagen density shows a very strong linear relationship with TDP-43 pathology. Scatterplot illustrating the strong linear relationship between skin collagen density index (CD-index) and TDP-43 pathology burden amongst skin biopsies (n=23 biopsies from 15 individuals) taken from the validation cohort. Skin collagen density explains 93% of the variation in TDP-43 pathology burden (p<0.001).

Of the 25 biopsies examined, 22 contained peripheral nerves and blood vessels, 15 contained sweat glands, and 12 contained sebaceous glands (Figure 7A). TDP-43 pathology was detected in 14/15 sweat gland-containing samples (93%), 7/12 sebaceous gland-containing samples (58%), 19/22 peripheral nerve-containing samples (86%), and 19/22 blood vessel-containing samples (86%) (Figure 7B). Across all of these structures, isolated cytoplasmic TDP-43 pathology was uncommon, observed in only 2/22 peripheral nerve, 5/15 sweat gland, 0/12 sebaceous gland and 3/22 blood vessel containing samples. Whilst nuclear-only pathology was also infrequent in peripheral nerves (2/22), blood vessels (4/22) and sweat glands (0/15), this was the most common pathology in sebaceous glands (6/7). The majority of peripheral nerves (17/22), sweat glands (9/15) and blood vessels (12/22) demonstrated both cytoplasmic and nuclear pathology. When stratified for severity level (mild, moderate and severe), peripheral nerves (n = 19 positive) most frequently showed severe pathology (9/19, 47%), with moderate (5/19, 26%) and mild (5/19, 26%) cases also represented. Blood vessels (n = 19 positive) displayed a similar distribution, with 9/19 (47%) severe, 6/19 (32%) moderate, and 4/19 (21%) mild. Sweat glands (n = 13 positive) were most often severely affected (10/13, 77%), with fewer cases showing mild (3/13, 23%) or moderate (1/13, 8%) pathology. In contrast, sebaceous glands (n = 7 positive) demonstrated a more balanced profile, with 4/7 (57%) severe and 3/7 (43%) mild, but no moderate cases.

#### Sweat and sebaceous glands show highest TDP-43 pathology

Among adnexal structures, sweat glands demonstrated the most consistent and robust TDP-43 pathology, with only one negative case, making them a suitable target for systematic quantification of pathology burden (Figure 7C). A superpixels-based segmentation was conducted across 15 samples from 11 patients, with H-scores calculated for 3–6 ROIs per sample. When grouped by anatomical site (Figure 7D), the greatest pathology burden was observed in the back (Average H-score 124.7, shoulder (Average H-score 121.1), and cheek (Average H-score 101.51), followed by arm (Average H-score 74.0), scalp (Average H-score 48.4), lip (Average H-score 42.0), neck (Average H-score 41.7), forearm (Average H-score 29.1), and chest (Average H-score 4.6) (Figure 7E). Pathology burden was significantly greater in sweat glands from sun-protected regions (arm, shoulder, back, Average H-score 104.2) compared with sun-exposed regions (forearm, lip, neck, scalp, cheek, Average H-score 50.8) (p=0.0037; Mann-Whitney U-test) (Figure 7F).

Furthermore, when sweat gland pathology was plotted against time to ALS diagnosis, heterogeneous trajectories were revealed (Figure 7G). In cases with multiple samples per patient, distinct temporal patterns emerged. For TR355_9, TDP-43 burden remained consistently high across both the arm (<1 year PSO, H-score 56.1) and neck (10.6 years PSO, H-score 47.5). In contrast, TR355_14 displayed striking regional variation: minimal pathology in the chest (1.5 years PSO, H-score 4.6) but markedly higher burden in shoulder (6.4 years PSO, H-score 125.7) and back (18.4 years prior, H-score 126.8). Regional comparisons across patients further confirmed this heterogeneity. Arm pathology was stable and high for TR355_11 (H-scores 85.7–86.2 at 5.2–5.3 PSO), but comparatively lower in TR355_9 near diagnosis (H-score 56.1 at 0.6 years PSO). Lip samples demonstrated greater burden closer to diagnosis (TR355_13, 5.2 years PSO, H-score 90.2) relative to lower pathology in samples obtained >10 years PSO (TR355_6 and TR355_8, H-scores 26.8 and 33.1). Conversely, shoulder (TR355_14 and TR355_19) and back (TR355_14 and TR355_15) regions consistently exhibited high levels (H-scores >116) irrespective of time to diagnosis.

#### Skin collagen density is strongly associated with TDP-43 pathology burden

To examine the relationship between skin collagen density and TDP-43 pathology burden, an average collagen density score (“Collagen density index” or CD-index) was calculated (along with average TDP-43 pathology burden as H-score, described previously) for each of 25 skin biopsies in the validation cohort. Average H-score and CD-index for each of the 25 biopsies is reported in Table 3, along with interval (in years) between skin biopsy sampling and diagnosis with ALS.

**Table 3.** Skin collagen density indices (CD-index) and TDP-43 pathology burden (H-score) score for skin biopsies in the validation cohorts. For each of the 25 skin biopsies in the validation cohort, collagen density indices (CD-index) and TDP-43 pathology burden (H-score) scores are reported, along with the interval (in years) between skin biopsy sample being taken an diagnosis with ALS.

| Table 3. Skin collagen density indices (CD-index) and TDP-43 pathology burden (H-score) scores for skin biopsies in the validation cohorts |  |  |  |
| --- | --- | --- | --- |
| Biopsy ID | TDP-43 pathology burden<br>H-score | Collagen density index<br>CD-index | Skin biopsy - ALS diagnosis interval (years) |
| TR355_1 | 91.82 | 1.53 | 6.32 |
| TR355_11A | 127.97 | 1.89 | 5.34 |
| TR355_11B | 184.26 | 3.01 | 10.42 |
| TR355_12<sup>†</sup> | 227.32 | 5.96 | 17.99 |
| TR355_13 | 85.09 | 1.49 | 0.58 |
| TR355_14A | 83.05 | 1.51 | 5.20 |
| TR355_14B | 77.25 | 1.47 | 5.24 |
| TR355_14C | 68.78 | 1.45 | 6.45 |
| TR355_14D | 90.50 | 1.54 | 6.71 |
| TR355_15 | 130.82 | 2.10 | 1.47 |
| TR355_16<sup>†</sup> | 278.40 | 11.03 | 18.50 |
| TR355_18 | 121.77 | 1.84 | 18.41 |
| TR355_19 | 150.66 | 2.39 | 23.52 |
| TR355_23 | 111.76 | 1.75 | 26.07 |
| TR355_2A | 148.99 | 2.20 | 6.32 |
| TR355_2B | 135.01 | 2.27 | 25.84 |
| TR355_3 | 137.12 | 2.08 | 17.98 |
| TR355_4A | 189.09 | 3.08 | 1.05 |
| TR355_5 | 145.27 | 2.32 | 6.69 |
| TR355_6 | 176.69 | 3.38 | 10.60 |
| TR355_8A | 158.60 | 2.56 | 26.57 |
| TR355_8B | 202.99 | 3.33 | 3.64 |
| TR355_8C | 160.03 | 2.45 | 23.08 |
| TR355_9A | 186.58 | 3.03 | 11.12 |
| TR355_9B | 122.44 | 1.88 | 11.27 |

<sup>†</sup> Biopsies TR355\_12 & TR355\_16 were identified as outliers with particularly high CD-index and/or H-score

Initial data inspection revealed two skin biopsies, both having particularly high H-score and CD-index, as data outliers - one biopsy taken one from back (TR355_16) and one from neck (TR355_12). For biopsy TR355_16, H-score and CD-index were identified as clear outliers (H score=278.40, CD-index=11.03; Table 3). For biopsy TR355_12, although CD-index was identified as an outlier (CD-index=5.96), H-score did lie within 1.5 times the interquartile range, whilst being the second highest (H score=227.32) of all biopsies, second only to TR355_16 (see above). Both (TR355_16 and TR355_12 were, therefore, excluded from these analyses, leaving 23 biopsies from 15 individuals. A very strong association between collagen density and TDP-43 pathology burden was uncovered (ρ=0.98, p<0.001; Spearman’s rank correlation), with collagen density explaining 93% of the variation in skin TDP-43 pathology levels (F_1,21_=277.40, R²adj.=0.93, β=60.68, p<0.001; Figure 7) in our sample of 23 skin biopsies. This general finding of association was robust to the inclusion of the two outlier biopsies TR355_16 and TR355_12 (F_1,23_=56.39, R²adj.=0.70, β=21.42, p<0.001), which is we report here (see also Supplementary Figure 3 scatterplot with outliers included) to illustrate that whilst there is a clear, strong relationship between collagen density and TDP-43 pathology burden across individuals, for a few individuals (2/17 here) particularly high skin collagen density does reflect high TDP-43 pathology too.

## DISCUSSION

For ALS, increased recognition of non-motor symptoms (*16, 60–62*), along with evolving evidence of pre-symptomatic and prodromal disease states (*63, 64*) are challenging traditional view of views of disease, with suggestions that adopting biological, rather than clinical, definitions of disease as with Huntington’s disease (*65*), Alzhemier’s disease (*66*), and neuronal α-synuclein disease (*67*), might be appropriate for diseases defined by TDP-43 proteinopathy (*68*). Biomarker evidence from pre-symptomatic stages of these diseases has been instrumental in reframing these traditional views, and here we present such evidence for ALS, i.e. evidence of pre-symptomatic disease-associated proteinopathy from a wide range of peripheral organ systems decades prior to motor symptoms, with skin TDP-43 pathology as a lead candidate biomarker for early detection of disease. Here, we discuss our findings in the context of past clinical translational failures, emerging views of ALS disease etiology, and future translational impact with real-life consequences for patients and people at risk of ALS.

Over the past two decades, more than 125 clinical trials have investigated 76 different drugs and enrolled over 15,000 individuals with amyotrophic lateral sclerosis (ALS). While our understanding of ALS pathobiology has advanced considerably during this time, these efforts have yielded limited translational success (*69*). This gap between biological insight and therapeutic impact is multifactorial, but one critical missing piece remains: the lack of reliable biomarkers capable of identifying at-risk individuals early in the disease course, before extensive neuronal loss has occurred. CNS-focused approaches to early diagnosis are not only invasive and impractical for point-of-care use, but also often detect changes too late, by which time up to 70% of motor neurons may already be lost.

In this context, recent findings highlighting systemic manifestations of ALS outside the CNS, such as in the skin (*39, 41, 48*) and gastrointestinal tract (*20–24, 26, 27, 29, 32, 48*), suggest that peripheral tissues may provide more accessible sites for diagnostic sampling. At the same time, recent advances in pathology detection tools such as the development of RNA aptamers for pathological TDP-43 (*57, 59*) and FUS (*70*), along with cryptic exon BaseScope^TM^ probes for *STMN-2* (*59*) and *HDGFL2* (*71*), are empowering fundamental new discoveries in ALS. Here, we employ two of these recently developed tools, TDP-43^apt^ in conjunction with *STMN-2* cryptic exon BaseScope^TM^ probe, to investigate address two crucial questions in ALS, i.e., (i) what is the extent of pre-symptomatic TDP-43 pathology detectable in an ante-mortem cohort of people who went on to develop ALS, and (ii) can we identify and validate a particular peripheral tissue as a biomarker with greatest potential for clinical applications to identify early TDP-43 pathology prior to disease symptom onset

In our current study, we identified TDP-43 pathology in skeletal muscle biopsies-specifically within the sarcoplasm and myonuclei of myocytes-features that were not previously discernible in the same cohort using earlier antibody-based approaches (*28*) (Figure 2A). Although other groups have demonstrated detection of TDP-43 pathology using pTDP-43 antibodies (*51–54*), this has only been from muscle from individuals who already received their diagnosis rather than pre-symptomatically as in the present study. Beyond confirming muscle involvement, we observed loss-of-function cryptic exon events in peripheral nerve bundles and vascular structures located between myocyte units. While TDP-43 misfolding within myocytes has been associated with physiological stressors such as hyperexcitability (*72, 73*), the presence of cryptic exon pathology in neurovascular bundles-outside the muscle units-points to a distinct and pathogenic process.

Furthermore, not only did we identify TDP-43 pathology in vascular and nerve structures in muscle tissue, but we also identified them in the skin, lymph nodes and GI tissue. These observations build on existing evidence of vascular TDP-43 pathology, including its presence in atherosclerotic plaques (*74*) and its association with comorbid arteriolosclerosis in the brain (*75*). Moreover, reduced nuclear TDP-43 in endothelial cells has been documented in postmortem ALS, frontotemporal dementia (FTD), and Alzheimer’s disease (AD) cortex (*50*). Loss of TDP-43 function in endothelial cells has also been shown to induce neuroinflammatory changes (*76*), compromise blood-brain barrier (BBB) integrity (*77*) and disrupt BBB-associated gene networks (*50*). This mirrors pathological mechanisms seen in other proteinopathies, such as beta-amyloid aggregation contributing to small vessel disease in AD (*78*). Whether TDP-43 aggregation in peripheral vasculature plays a similar systemic role remains to be fully elucidated. However, its detectability in end-organ blood vessels represents a promising opportunity to develop minimally invasive, blood- and/or biopsy-based diagnostic biomarkers that could enable earlier identification of individuals at risk for ALS.

We also observed more consistently detectable TDP-43 pathology in skin (n=7/8 biopsies from a total of 7 patients; previously 2/8 biopsies from 7 patients) biopsies using the TDP-43 aptamer, compared to antibody-based approaches (*28*). In one case of note, an individual with a *C9orf72* mutation had two temporally distinct biopsies from the same skin site. The first biopsy (a diagnostic excision biopsy of skin from the chest, containing a squamous cell carcinoma *in situ*) was taken 24 months prior to symptom onset. Within 12 months of diagnosis with ALS - a second biopsy (a further diagnostic excision biopsy, containing a benign squamoproliferative lesion) was taken from the same anatomical site as the first. We observed no pathological TDP-43 in the first biopsy but found florid TDP-43 pathology in the second biopsy. Whilst this is only a single case, it demonstrates the potential of skin as a phenoconversion time-window for monitoring the emergence of TDP-43 pathology. Conversely, in our validation cohort, with a larger sample size, TDP-43 pathology was seen to be more temporally robust, highlighting that more research is required before skin could be usefully deployed in the context of tracking phenoconversion, and might be better placed as a tool for trial stratification or for target engagement. Indeed, skin pathology might provide insights in pathological protein aggregation beyond TDP-43, as indicated by a recent case-study instance of FUS protein nuclear loss detected from the a skin biopsy of an individual who presented with cramps and progressive limb weakness over a 2-year period, with a family history of progressive neuromuscular deterioration, whose FUS-ALS diagnosis was confirmed through the identification of a FUS-ALS associated (c.1562G>T; p. R521H) mutation (*79*).

In our validation cohort, we examined skin samples obtained up to 26.5 years prior to symptom onset, encompassing a diverse range of anatomical sites, including the skin of the head and neck, trunk, perineum, and both upper and lower limbs. Remarkably, pathological TDP-43 was detectable across all sampled skin sites and very strongly associated with structural changes, reinforcing the notion of widespread peripheral involvement years before any signs or symptoms. The localisation of pathology to sweat and sebaceous glands is particularly noteworthy, as these structures are innervated by peripheral autonomic fibers. Their involvement suggests that pathological TDP-43 accumulation may not be limited to motor neurons but could extend to broader components of the peripheral nervous system, including autonomic pathways. Similar to the proposed propagation of α-synuclein via the vagus nerve in Parkinson’s disease (*80–82*), TDP-43 pathology may also originate or spread via peripheral-autonomic routes. Understanding the ontogeny of this pathology will be crucial for refining early detection strategies. Moreover, the presence of TDP-43 pathology within the sebaceous glands is reminiscent of individuals with Parkinson’s disease, where pre-symptomatic α -synuclein pathology can be detected from sebum (*83*), and lipidomic analysis of skin swabs can differentiate people with Parkinson’s disease, Alzheimer’s disease, and healthy subjects (*84*).

The consistent detection of TDP-43 pathology in these glandular structures, particularly sweat glands, also presents a practical advantage: they are among the most accessible targets for minimally invasive skin biopsies and hold potential for future liquid biopsy applications. In our cohort, the majority of sweat glands examined were positive for pathology, further supporting their robustness as a biomarker source. If validated in larger and longitudinal cohorts, including individuals at ALS risk prior to phenoconversion, sweat gland pathology could form the basis of a scalable and repeatable peripheral assay for early ALS detection. Such an approach would address a critical gap in the field by enabling earlier diagnosis, stratification for clinical trials, and monitoring of disease progression-without reliance on invasive CNS sampling, or late-stage symptom emergence.

Collectively, these findings represent a paradigm shift in ALS biomarker research. By demonstrating that TDP-43 pathology can be robustly detected in peripheral tissues, including skin, muscle, lymph nodes, and gastrointestinal organs, years before symptom onset, our work highlights the potential of accessible, minimally invasive tissues as windows into pre-symptomatic disease biology. Skin, in particular, emerges as a highly practical and informative substrate, with back and shoulders providing reproducible anatomical sites for early pathological detection. Furthermore, the identification of reproducible and temporally stable pathology in the sweat and sebaceous glands at these sites highlights potential for non-invasive skin surface detection. This peripheral focus addresses a critical gap left by CNS-centered biomarker discovery approaches, which are invasive and typically capture disease only after substantial neuronal loss. The ability to detect TDP-43 pathology pre-symptomatically not only offers a path toward earlier and more equitable diagnosis but also provides a platform for monitoring disease progression, evaluating therapeutic interventions, and stratifying participants for clinical trials. In doing so, peripheral tissue biomarkers fundamentally transform the ALS field, moving it toward a model in which disease can be identified and potentially targeted long before motor symptoms appear, paralleling the advances seen with peripheral α-synuclein detection in Parkinson’s disease.

Our findings highlight the potential utility of skin TDP-43 pathology as an early, accessible biomarker for trial stratification for example in FTD, where underlying molecular subtypes such as FTD-TDP, FTD-TAU, and FTD-FUS have distinct prognostic and therapeutic implications. Detecting TDP-43 misfolding in peripheral tissues could allow earlier identification of individuals most likely to benefit from TDP-43-targeted interventions, while excluding those with non-TDP-43 pathologies from trials where they are unlikely to respond. Beyond FTD, skin TDP-43 pathology could hold promise across the spectrum of TDP-43 proteinopathies. In primary TDP-43 proteinopathies (including FTD, amyotrophic lateral sclerosis, frontotemporal lobar degeneration, and disorders such as Perry syndrome and FOSMN) early peripheral detection could refine diagnosis, prognosis, and patient selection. In addition, the presence of secondary TDP-43 pathology in diseases such as AD, chronic traumatic encephalopathy, and PD suggests broader utility in mapping disease heterogeneity and identifying patients at risk of mixed or evolving proteinopathies. Together, these observations position skin TDP-43 pathology as a scalable biomarker platform for precision medicine approaches across diverse neurodegenerative disorders.

### Study limitations

Despite the compelling evidence for widespread peripheral TDP-43 pathology in ALS, several limitations should be acknowledged. First, the sample size, particularly for ante-mortem tissue cohorts, remains modest, limiting the statistical power to fully assess the variability of peripheral pathology across different ALS subtypes and genetic backgrounds. Second, while our study identifies TDP-43 pathology in multiple peripheral organs, the observational design precludes definitive conclusions regarding causality or the temporal sequence of pathological spread between central and peripheral tissues. Third, the validation of skin as a biomarker substrate, although promising, requires further longitudinal studies with repeated biopsies and correlation with clinical outcomes to determine sensitivity, specificity, and predictive value across diverse populations. Fourth, while our analyses focused on TDP-43 gain- and loss-of-function pathology, other molecular or cellular changes in peripheral tissues may contribute to disease pathogenesis but were not assessed in this study, for example the effects of the immune system. Finally, tissue accessibility and preservation could introduce sampling bias; for example, ante-mortem skin biopsies may not fully represent the systemic burden of TDP-43 pathology. Despite these limitations, our findings provide a strong rationale for the continued exploration of peripheral tissues as a minimally invasive biomarker platform and a window into the early pathobiology of ALS.

## MATERIALS AND METHODS

### Tissue provision and ethics

Tissue for the biomarker discovery cohort was requested from the Lothian NRS BioResource RTB ethical approval 15/ES/0094, covering use of residual tissue surplus to diagnostic requirements taken as standard of care with approval number SR1684. Tissue for the biomarker validation cohort was requested from NHS Grampian Biorepository RTB ethical approval 21/NS/0047 with approval number TR355. All archived formalin-fixed, paraffin-embedded (FFPE) tissue material with an NHS diagnostic code (e.g. any tissue samples taken for diagnostic or surgical purposes at any time during the patient’s life, under the care of NHS Lothian and NHS Grampian) were requested for all patients with known diagnosis of ALS. The tissue was then assessed for sufficiency by two independent histopathologists (SBP & JMG) using a single H&E. No identifiable clinical or demographic data were used in this study. Tissue provided by the BioResources had been fixed in 10% formalin for a minimum of 72 h, then dehydrated in an ascending alcohol series (70–100%) followed by three 4-hour washes in xylene. Three successive 5-hour paraffin wax embedding stages were performed followed by cooling and sectioning of tissue on a Leica microtome into 4 μm thick serial sections that were collected on Superfrost (ThermoFisher Scientific) microscope slides. Sections were dried overnight at 40°C before staining. Bleach treatment was performed (on skin cases where warranted) by incubating slides immediately post antigen retrieval in 10% hydrogen peroxide for 12 hours overnight. This step was followed by the normal protocol in full, including the additional 3% hydrogen peroxide step used to block endogenous peroxidase activity, following a previously published and optimised protocol with SOP (*85*).

### Immunohistochemistry and BaseScope^TM^ in situ hybridisation

Sections were dewaxed using successive xylene washes followed by alcohol hydration and treatment with picric acid for removal of formalin pigment. For TDP-43^APT^ staining, antigen retrieval was carried out in citric acid buffer (pH 6) in a pressure cooker for 15 min, after which immunostaining was as described previously (*14, 58*). *In situ* hybridisation was performed on tissue sections using BaseScope^TM^ reagents (Advanced Cell Diagnostics) as per the manufacturer’s instructions (*86*) and adapted as described previously (*87*). Probe hybridisation was performed using BaseScope^TM^ probes for *STMN-2* CE mRNA transcripts (catalogue number 1048231-C1). Slides were counterstained using haematoxylin and lithium carbonate, washed in xylene, and coverslips were mounted using DPX Mountant. Slides were imaged using an EVOS M5000 microscope using brightfield settings at 20x and 40x magnification.

### Imaging and digital pathology analysis

Brightfield photomicrographs were taken at 20X for digital pathology analysis and 40X for capturing high resolution images using an EVOS M5000 microscope. For digital pathology analysis of sweat glands, three 50×50 µm regions of interest (ROIs) were chosen from each photomicrograph using QuPath (*88*). In brief, mean DAB intensity for each case stained with TDP-43 RNA aptamer was digitally acquired by superpixel analysis using digital burden scoring performed using the freely available QuPath software implementing superpixel analysis using code published previously (*89, 90*). The mean DAB intensity of all sweat gland ROIs was calculated and exported for subsequent analysis. Difference of Gaussian (DoG) superpixel segmentation analysis was then carried out on the sweat glands to quantify aptamer-positive foci. Here, compartments were split into superpixels that were generated from pixels with similar intensities and textures for further classification. Each superpixel was classified as negative or varying levels of positive, depending on the pre-set DAB intensity thresholds. The H-score from each ROI was then exported. Then, H-scores were calculated first per case by averaging the 3 ROIs from each sample, and then per region by averaging the H-score from each case.

### Statistical analysis

Statistical analyses and plot visualisation were carried out using GraphPad Prism (version 10) and R (*91*) (version 4-5.1). TDP-43 pathology burden scores (H-score) and collagen density indices (CD-index) were averaged over three ROIs, when available, for each sample. Statistical comparisons were only conducted between groups with *n* ≥ 3. The normality of continuous variables was assessed using the Shapiro–Wilk test (*shapiro.test* function, *stats* (*91*) package). Homogeneity of variances was evaluated using Levene’s test (*leveneTest* function, *car* (*92*) package). Outliers were identified as those 1.5x outside the interquartile range. Where appropriate, non-parametric Mann-Whitney U statistical test was used to compare two independent groups. Conservatively, Spearman’s rank correlations were used to assess associations between continuous variables, and linear regression models were fitted using R’s *lm*() function. Interval periods (in years) between date of biopsy and date of ALS diagnosis were calculated by dividing the elapsed time between dates, in days, by 365.25. The *tidyverse* (*93*) package was used for data manipulation and visualisation; specifically, data summaries were computed using *dply* (*94*) summary functions (and base R (*91*)), data were visualised and plotted using *ggplot2* (*94*).

## Author contributions

*JMG* conceptualized, secured primary funding for, and led the study with key input from *FMW*, *KH*. Patients for the discovery cohort were identified by *SBP*, *JMG* with tissue request prepared and coordinated by *JMG*. Patients for the validation cohort were identified by *ADM* with tissue request prepared and coordinated by *KR*. Immunohistochemistry was performed by *KH*, *AR*, *FMW*, *TL*, and *FLR*. *KH*, *AR*, and *JMG* undertook digital image acquisition and curation. Data curation was performed by *KH*, *FMW*, *AR* and *JMG*. Data analysis was carried out by *FMW*, *KH*, *JMG*. Early drafts were prepared by *FMW*, *JMG*, with input from *KH*, *HS*, *KR*, *SBP*. *FMW* drafted the initial manuscript with input from *KH*, *JMG*. Subsequent drafts were prepared by all authors and finalized by *FMW*, *KH* and *JMG*.

## Competing interests

Authors declare that they have no competing interests.

## Data and materials availability

All data are available in the main text or the supplementary materials.

## Funding

Funders had no role in study design, data collection, data analyses, interpretation, or writing the manuscript. The research leading to this manuscript has been supported by: (i) a Target ALS Early-Stage Clinician Scientist award to *JMG* (FS-2023-ESC-S2), (ii) an NHS Grampian grant to *FMW* (GCA25107); (iii) an NIH grant to *JMG* and employing *HS* and *FR* (R01NS127186); a (iv) MND Scotland/CSO grant to *KR* (CAF/24/13 – 2024/MNDS/6500/780ROB); (v) a Motor Neurone Disease Association grant to *JMG* and employing *KH* (Gregory/APR24/2374-791); (vi) a LifeArc MND Primer Fund to *JMG* (A20170). Funders had no role in study design, data collection, data analyses, interpretation, or writing the manuscript. The authors would also like to thank the University of Aberdeen Microscopy and Histology Core Facility in the Institute of Medical Sciences, and the NHS Lothian BioResource staff, in particular Vishad Patel and Craig Marshall.

## Acknowledgements

The authors would also like to thank the University of Aberdeen Microscopy and Histology Core Facility in the Institute of Medical Sciences and the NHS Lothian BioResource staff, in particular Vishad Patel and Craig Marshall.

---

**Supplementary Figure 1.**
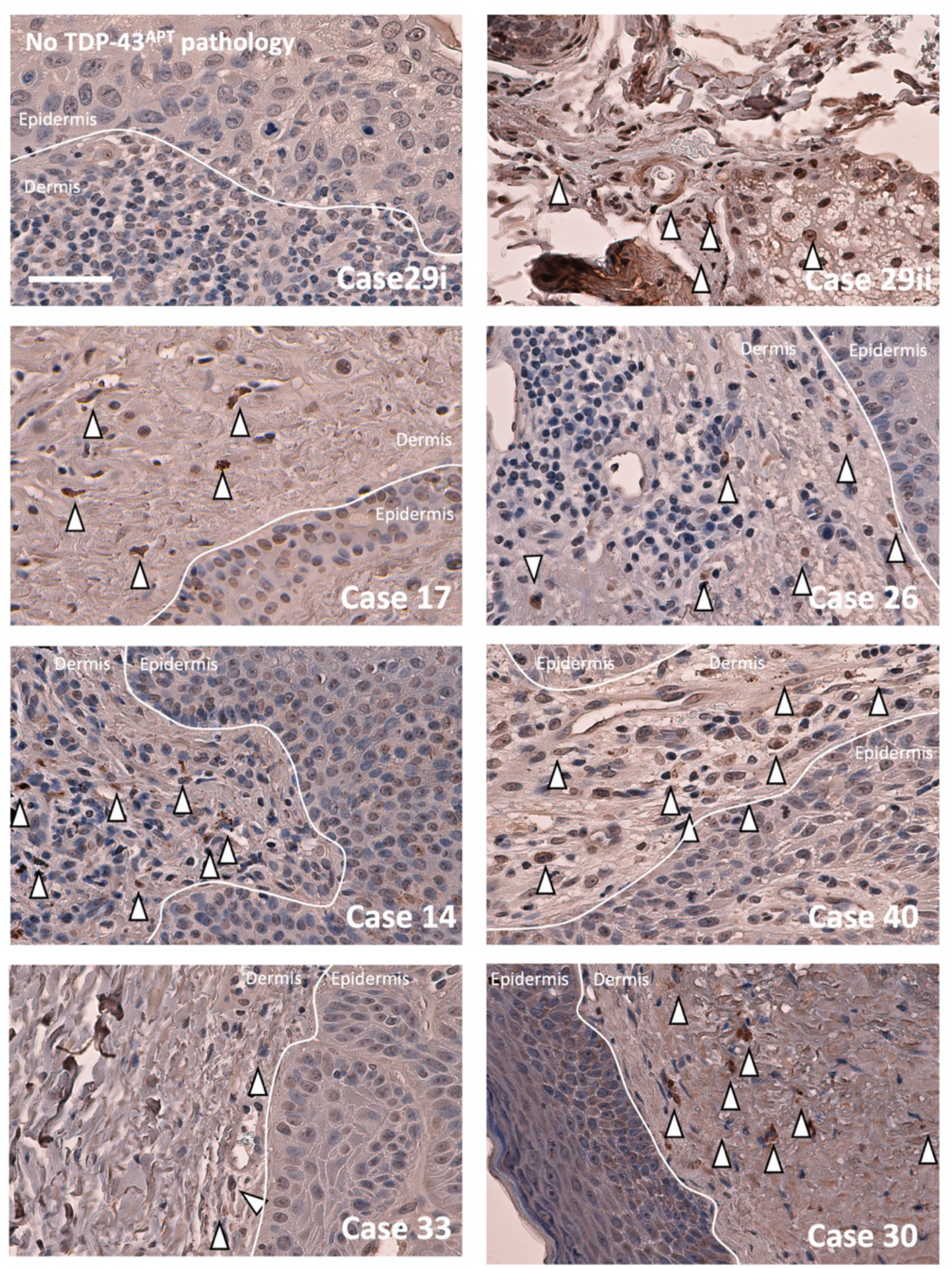
TDP-43^APT^ pathology detected in all skin biopsies from individuals who went on to develop ALS. Photomicrographs taken at 40x of all skin biopsies from this cohort (n=7 individuals). One individual (case 29), who had a *C9orf72* mutation, had two biopsies taken 24 months apart (29i and 29ii). Sections are stained with TDP-43^APT^ (DAB chromogen in brown) and counterstained with haematoxylin (blue). TDP-43^APT^ pathology is indicated with white arrowheads and can be seen within the dendritic cells of the dermis, within neurovascular bundles, and sebaceous units. Delineation between dermis and epidermis is indicated by a continuous white line. Patient ID (case number) relates to clinical demographics published previously for this cohort (*28*). Scale bar = 20 μm.

**Supplementary Figure 2.**
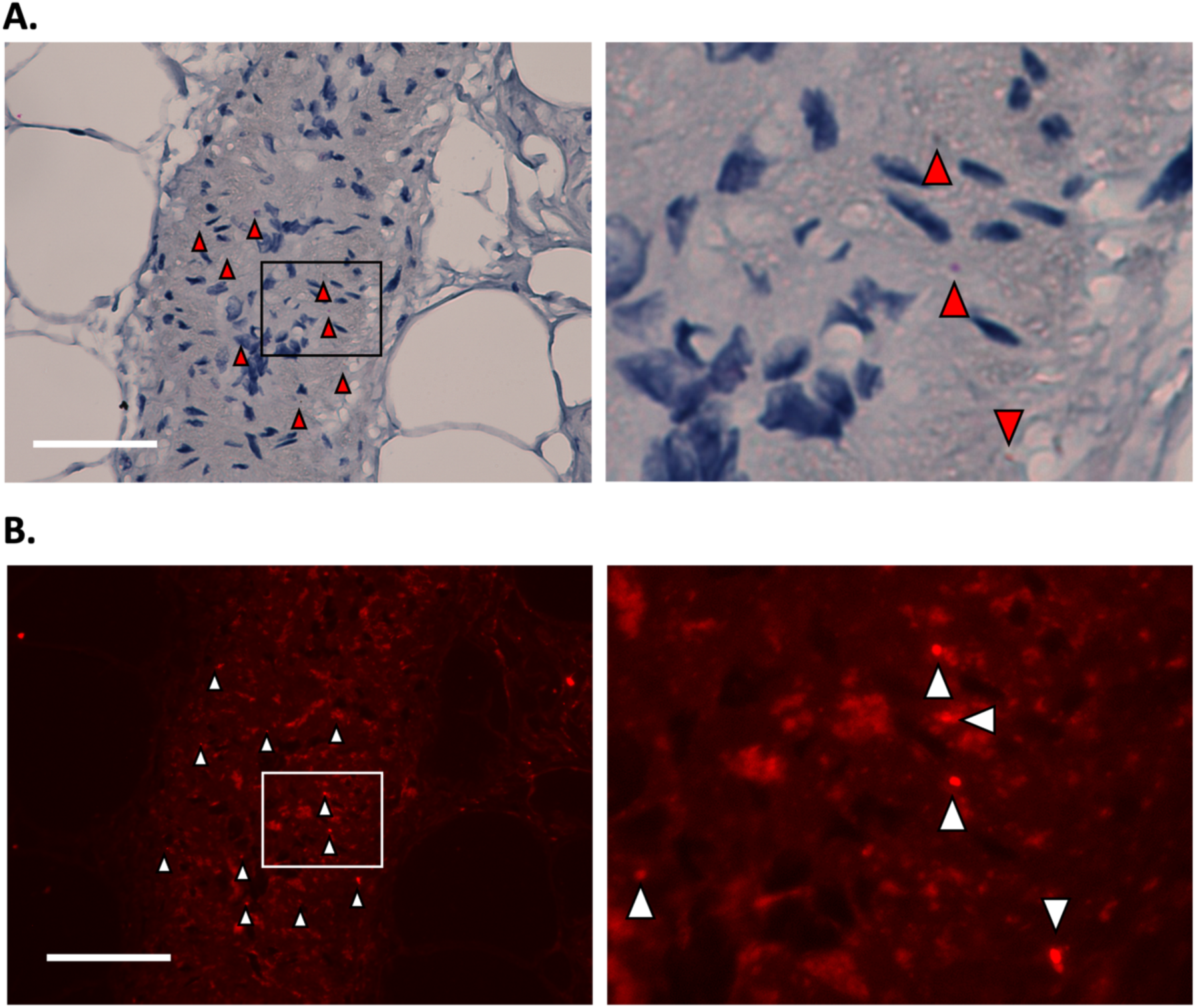
TDP-43 pathology seen in neurovascular bundle within the fibroadipose tissue separating fascicles of muscle fibres. **A.** Photomicrographs taken at 20x magnification (left image), and 4 times optical zoom of the area highlighted with the black box (right image) stained with *in situ* hybridisation probes directed against the cryptic exon of STMN-2 and counterstained with haematoxylin. Signal can only be seen in the context of a loss of TDP-43 function, where it is no longer repressing cryptic exon inclusion. **B**. The same frame from (**A**) imaged using fluorescence microscopy (fast red chromogen is also a fluorophore), highlighting individual mRNA molecules of STMN-2 transcripts that contain cryptic exons indicating TDP-43 loss-of-function. Scale bar = 50 μm.

**Supplementary Figure 3.**
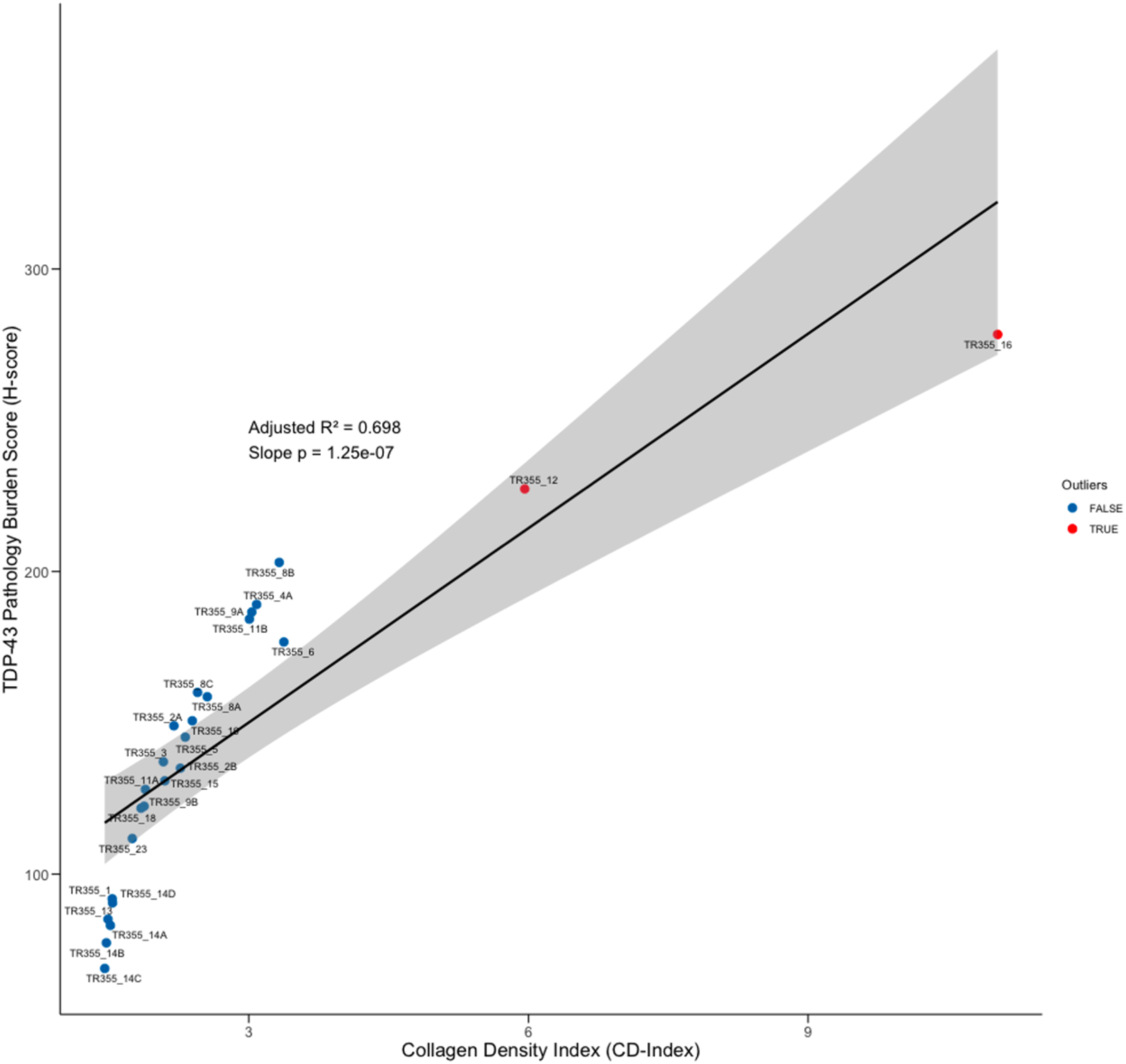
Skin collagen density shows a very strong linear relationship with TDP-43 pathology, robust to the inclusion of outlier samples TR355_12 and TR355_16. Scatterplot illustrating the strong linear relationship between skin collagen density index (CD-index) and TDP-43 pathology burden amongst skin biopsies when two outlier samples are also included (giving n=25 biopsies from 17 individuals) taken from the validation cohort.

**Supplementary Table 1.**
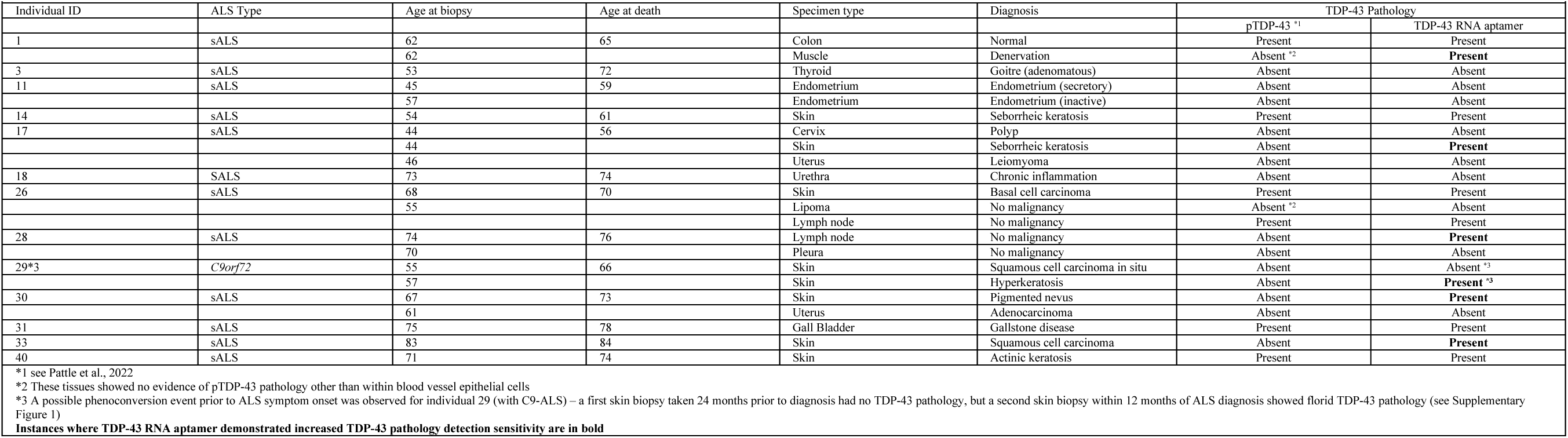
TDP-43 aptamer and STMN-2 cryptic exon BaseScopeTM in situ hybridisation probe pathology detection modalities provide greater sensitivity and equivalent specificity to antibody approaches for detecting TDP-43 pathology in ALS pre-symptomatic peripheral tissues. Summary table of clinicopathological information for the biomarker discovery cohort (n = 13 individuals) which was screened for TDP-43 pathology in this study, presented along with presence or absence of TDP-43 pathology detected previously using antibody approaches (*28*). Each row lists the individual ID, ALS subtype, age at biopsy, age at death, biopsy specimen type, contemporaneous pathological diagnosis, along with the presence or absence of TDP-43 pathology. For several individuals, multiple tissue samples collected at different times or from distinct anatomical sites are presented.

## Notes

### Competing Interest Statement

The authors have declared no competing interest.

### Summary of Updates

In this updated version, the authors have assembled and explored an additional validation cohort from a second referal centre, strengthening the findings of the original preprint. Our conclusions highlight the importance of skin TDP-43 pathology as an early biomarker of ALS.

